# The immunomodulatory effect of IrSPI, a tick salivary gland serine protease inhibitor involved in *Ixodes ricinus* tick feeding

**DOI:** 10.1101/705921

**Authors:** A. A. Blisnick, L. Šimo, C. Grillon, F. Fasani, S Brûlé, B. Le Bonniec, E. Prina, M. Marsot, A Relmy, S Blaise-Boisseau, J. Richardson, S.I. Bonnet

## Abstract

Ticks are strict hematophagous arthropods and are the most important vectors of pathogens affecting both domestic and wild animals worldwide. Moreover, they are second only to mosquitoes as vectors of human pathogens. Hard tick feeding is a slow process—taking up to several days for repletion prior to detachment—and necessitates extended control over the host response. The success of the feeding process depends upon injection of saliva by tick, which not only controls host haemostasis and wound healing, but also subverts the host immune response to avoid tick rejection during this long-lasting process. In turn, the manipulation of the host immune response creates a favourable niche for the survival and propagation of diverse tick-borne pathogens transmitted during feeding. Here, we report on the molecular and biochemical features and functions of IrSPI, an *Ixodes ricinus* salivary serine protease inhibitor involved in blood meal acquisition. Our results show that IrSPI harbours the typical conformational fold of Kunitz type I serine protease inhibitors and that it functionally inhibits the elastase and, to a lesser extent, chymotrypsin. We also show that IrSPI is injected into the host during feeding. Crucially, we found that IrSPI has no impact on tissue factor pathway-induced coagulation, fibrinolysis, apoptosis, or angiogenesis, but a strong effect on immune cells. IrSPI affects antigen-presenting macrophages by hampering IL-5 production. In addition, IrSPI represses proliferation of mitogen-stimulated CD4^+^ cells. The inhibition of T cell proliferation was associated with marked reductions in pro-inflammatory cytokine secretion. Our study contributes valuable knowledge to tick-host interactions and provides insights that could be further exploited to design anti-tick vaccines targeting this immunomodulator implicated in successful *I. ricinus* tick feeding.

**Author summary:** Ticks are the most important vector influencing both human and animal health in Europe, where *Ixodes ricinus* is the most abundant tick species. Ticks feed on animal or human blood for an extended period, during which their saliva allows both feeding and pathogen transmission by interfering with native host responses. A better understanding of tick-host-pathogen interactions is central to the discovery of improved control methods. Within this context, we previously identified IrSPI as an *I. ricinus* salivary molecule implicated in both tick feeding and bacterial transmission. This serine protease inhibitor was almost characterised as an elastase inhibitor, and here, we show IrSPI overexpression in several tick organs—especially salivary glands—during blood feeding. We demonstrate that IrSPI is injected into the host through saliva, and despite having no impact on endothelial cell angiogenesis or apoptosis during blood feeding, we report an immunomodulatory role, whereby CD4^+^ T lymphocyte proliferation is repressed and where the cytokine secretion pattern of both splenocytes and macrophages is modified. Our study provides new insights into the complex armament developed by ticks to overcome the host response, and uncovers a potential vaccine target for disruption of feeding processes and pathogen transmission.

## Introduction

Ticks are obligate hematophagous arthropods able to feed on diverse vertebrate hosts including mammals, birds, amphibians, and reptiles. They cause substantial economic losses in the livestock industry due to blood spoiling and secondary infection of bite wounds, which consequently decrease food production and the value of leather. Crucially, ticks are excellent vectors for pathogens, including bacteria, parasites, and viruses. They are—after mosquitoes—the second-most common vector of human pathogens, and the first for animal health [1]. In particular, the hard tick *Ixodes ricinus* is the most widespread and abundant tick species in Europe, as well as the most efficient transmitter of pathogens that significantly impact both human and animal health [2]. This tick species follows a three-host life cycle where each of the three distinct life stages (larvae, nymph and adult) feeds only once on its respective host.

The hard tick blood meal’ is one of the longest feeding processes of all blood feeding arthropods. Several hard tick species feed over ten days as adults, thus causing severe host tissue damage. The success of the blood meal is critically dependent on saliva, which represents the primary tick-host interface (reviewed in [3]). Indeed, tick saliva possesses several essential properties that facilitate feeding, including an ability to repress host pain at the tick bite site, thus preventing early detection by the host. In addition, tick saliva has the capacity to counteract host haemostatic and immunological responses at the site of injury. Ticks are pool-feeders, meaning that they create a haemorrhagic blood pool by alternating blood absorption and saliva injection. As such, the saliva’s anti-coagulant properties maintain the blood pool in a fluid state by blocking blood coagulation cascade activation and promoting fibrinolysis. Moreover, tick saliva impedes the host’s innate immune system, by preventing complement activation and immune cell recruitment, thus precluding tissue remodelling and angiogenesis as well as local inflammation. Tick saliva also impairs anti-tick adaptive immune responses, by repressing lymphocyte recruitment at the tick bite site. Finally, these tick bite-associated processes create an environment favouring the transmission of tick-borne pathogens (TBP) to the host.

A better understanding of tick-host-pathogen interactions is central to the development of improved tick and tick-borne disease (TBD) control methods [4]. In order to identify tick factors involved in both tick biology and pathogen transmission, we used next generation sequencing (NGS) to compare gene expression in salivary gland (SG) tissue from uninfected and *Bartonella henselae*-infected *I. ricinus* ticks [5]. This gram-negative bacterium is responsible for cat scratch disease in humans and is transmitted from cat to cat by fleas. Although the majority of human cases are due to scratches and bites from infected cats, we have previously demonstrated that *I. ricinus* can also directly transmit this pathogen [6]. A small percentage of tick genes (5.6%) were found to be modulated by this bacterial infection. The most overexpressed gene—that we named *Ixodes ricinus* serine protease inhibitor (*IrSPI*)—encoded a putative serine protease inhibitor. Preliminary RNA interference (RNAi) functional analysis revealed that IrSPI is implicated in tick feeding as well as in *B. henselae* infection of tick SG, rendering it a promising target as an anti-*I. ricinus* vaccine. It is also known that several serine protease inhibitors (SPI) are involved in the aforementioned tick saliva processes that maintain blood pools, further justifying our interest in IrSPI [7]. In fact, serine protease inhibitors are one of the main protein families injected into the host by ticks during blood feeding, as the proteins they inhibit play key roles in regulating host coagulation, angiogenesis, haemostasis, and immune responses. Moreover, in ticks, several serine protease inhibitors are involved in blood digestion, the tick’s innate immune system, tick reproduction, as well as pathogen transmission [7].

The aim of this study was to elucidate the role of IrSPI both in tick biology and at the interface of the tick and the vertebrate host during feeding. *In silico* characterization of IrSPI confirmed that it belonged to the Kunitz type I family of serine protease inhibitors. We showed that *IrSPI* was induced during blood feeding and that the protein was injected into the host, suggesting involvement in modulating host responses to tick feeding. We found that elastase was inhibited by recombinant IrSPI, whereas neither the tissue factor pathway blood coagulation, fibrinolysis, endothelial cell apoptosis nor angiogenesis were affected. Finally, we demonstrated that IrSPI modulates the host immune response by reducing the proliferative capacity of T lymphocytes and by altering the cytokine expression profile of both macrophages and splenocytes.

## Results

### IrSPI molecular characterisation

The full-length *IrSPI* cDNA encompasses a 321-nucleotide open reading frame encoding a 19-residue signal peptide, followed by a 285-nucleotide sequence encoding a mature 94 amino acid protein. *I. ricinus* and *I. scapularis* sequences presenting the greatest degree of IrSPI identity are shown in Fig 1A. The IrSPI sequence harbours seven cysteines which potentially form three disulfide bonds. The putative mature protein has a molecular mass of 8310 Da, a single potential N-glycosylation site, and ten putative phosphorylation sites (Fig 1B). The intact molecular mass of the recombinant IrSPI expressed in Drosophila S2 cells was analysed by mass spectrometry, using the UltrafleXtreme MALDI-TOF/TOF instrument (Bruker-Daltonics, Germany). Results revealed a 990 Da shift between the experimental mass (13,034 Da) and the theoretical mass (12,044 Da) of the recombinant protein, which is compatible with the addition of ten phosphoryl groups (80 Da) and a single N-linked glycan (161 Da) as previously hypothesised via NetNGlyc 1.0 and NetPhos 3.1 servers (Fig S1).

**Fig 1.**
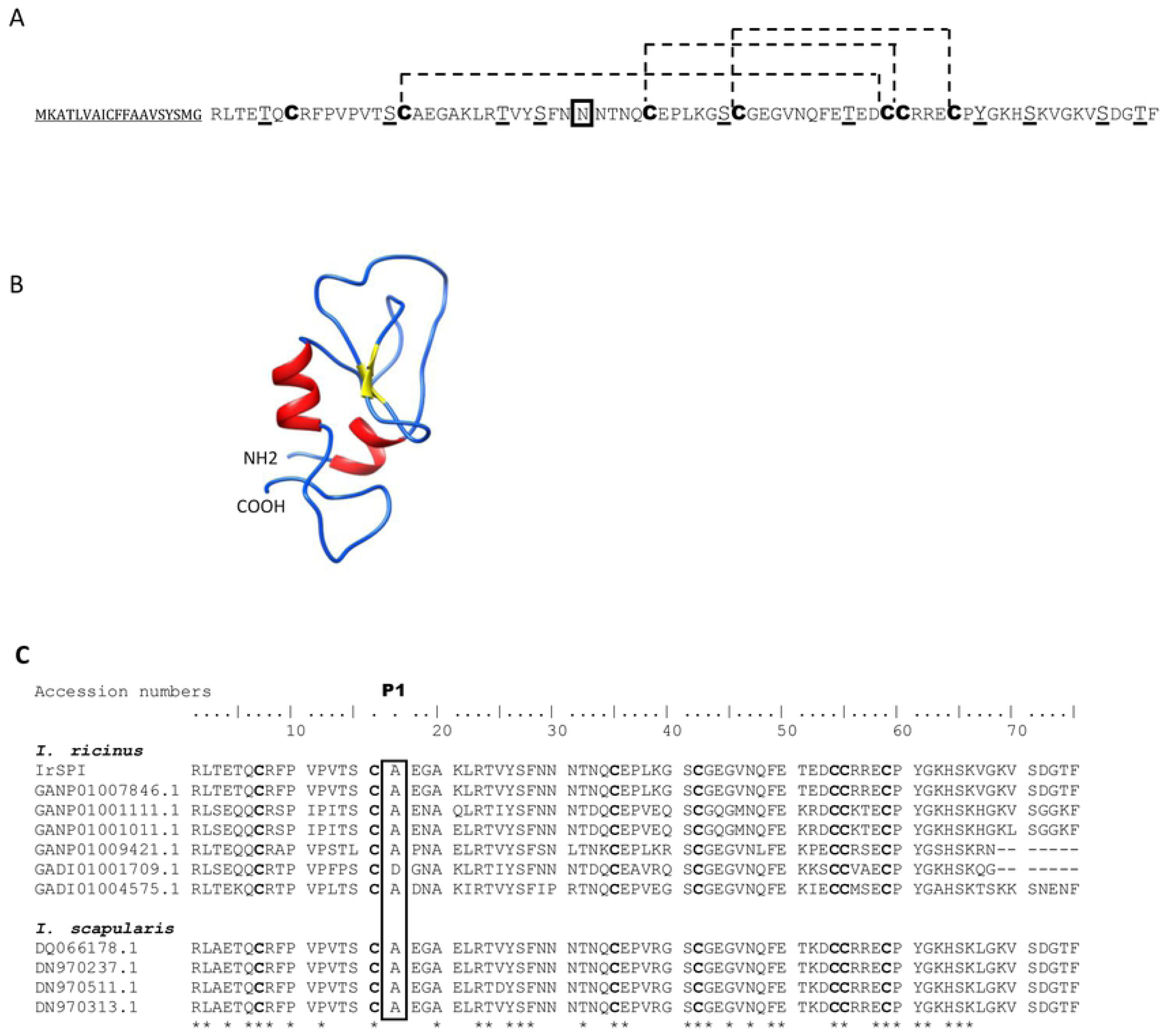
Structures of IrSPI (KF 531922.2) and amino acid sequence alignment of *I. ricinus* and *I. scapularis* Kunitz protease inhibitors. A) Primary and B) secondary IrSPI structures. The signal peptide is underlined. Post-translational modifications are predicted to include 10 putative phosphorylation sites (underlined) and 1 putative N-glycosylation site (double underlined). The 7 cysteine residues (in bold) are capable of forming 3 disulphide bridges (dashed line) and, combined with the formation of 2 β–sheets (yellow) and 2 α-helices (red), bestowing to the IrSPI protein its unique Kunitz/BPTI fold. C) Amino acid sequence alignment of IrSPI with other known *I. ricinus* and *I. scapularis* Kunitz protease inhibitors with corresponding accession numbers. Identical amino acids are marked with asterisks and the P1 residue is highlighted by a rectangle. Accession numbers in GenBank are indicated for each gene, as well as the percentage of identity with IrSPI.

The predicted protein structure, as determined *via* Phyre 2 software, consists of two β-sheets, two α-helices, and a single Kunitz-like inhibitory domain, suggesting that IrSPI belongs to the Kunitz type I protease inhibitor family (Fig 1C). Recombinant IrSPI protein was used to confirm this structure but circular dichroism experiments failed to reveal the expected α-helix signals (Fig S2). In addition, the main population observed *via* analytical ultracentrifugation had a sedimentation coefficient of 1.6S and a frictional ratio (f/f0) of 1.2, consistent with a monomeric and globular form of IrSPI (Fig S3). The absence of helices in the recombinant IrSPI structure may prevent appropriate disulfide bond formation, consequently causing hydrophobic residues to be exposed at the protein surface. Thus, the globular aspect of IrSPI could be due to an abortive folding process and its globular form might represent a folding intermediary, and not the correct functional IrSPI structure. Altogether, results suggest that the recombinant protein may be misfolded, which might compromise its functional activity as a protease inhibitor. Consequently, serine protease inhibition assays were performed with a refolded form of recombinant IrSPI and the refolding buffer was used as a control.

### Serine protease inhibition

Since the P1 residue of IrSPI is an alanine, thus we hypothesised that it could inhibit serine proteases of the elastase-like family. We thus assessed the ability of IrSPI to inhibit cleavage of the respective substrates of several serine proteases, including elastase. Although the non-refolded recombinant protein did not inhibit any serine protease; refolded IrSPI in excess (1 µM) moderately but significantly decreased elastase catalytic activity (23.9%; ANCOVA test, P=0.0002), and that of chymotrypsin very slightly (1.7%) (Fig 2). Trypsin, thrombin, and kallikrein were not inhibited.

**Fig 2.**
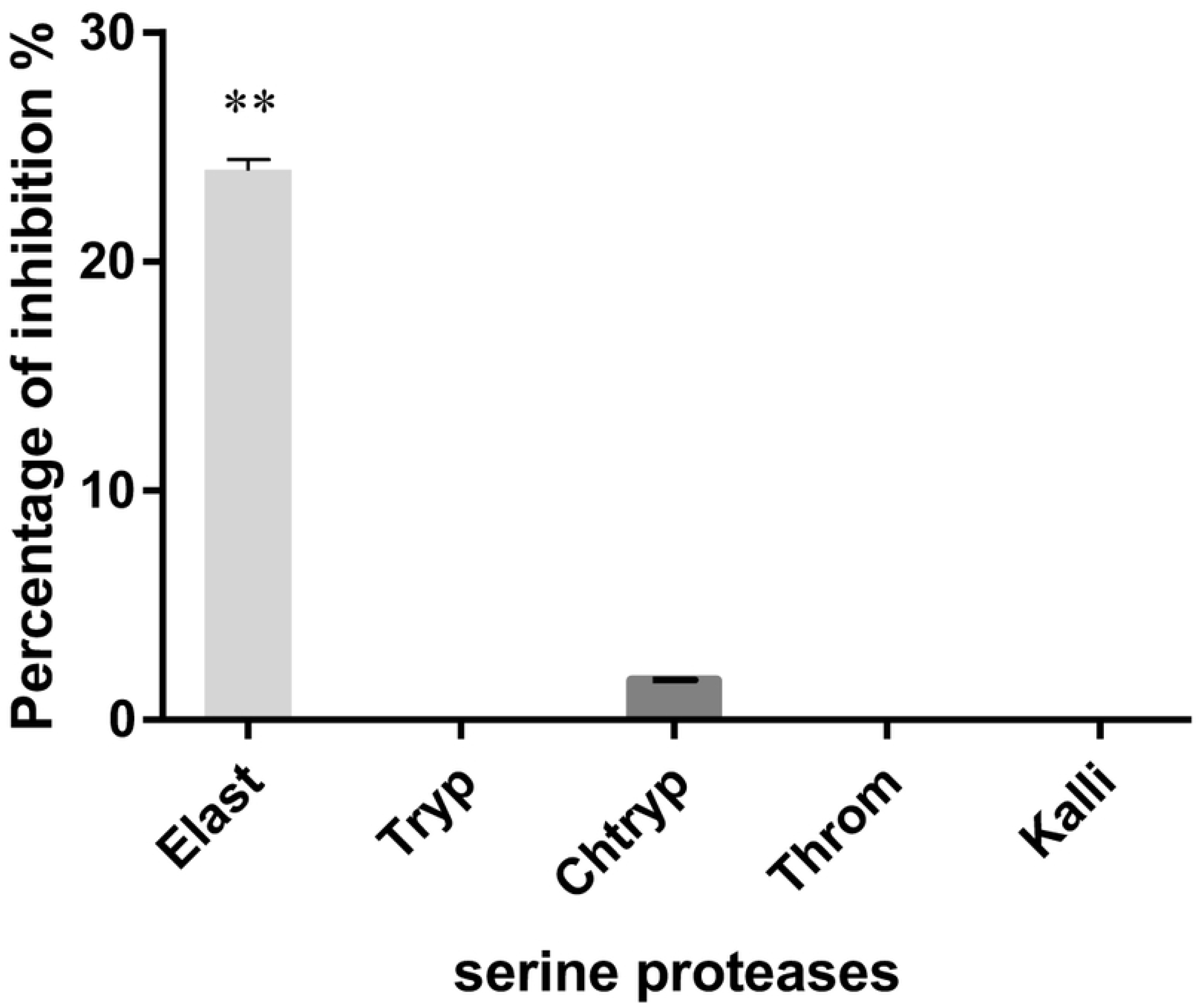
Inhibition of serine proteases by IrSPI. The ability of IrSPI (1µM of refolded protein) to inhibit the activity of elastase (Elast), trypsin (Tryp), chymotrypsin (Chtryp), thrombin (Throm), and kallikrein (Kalli) on synthetic substrates was evaluated at 37°C for 30 min. Enzyme activity was evaluated from the slope of A_405_ versus time and results were expressed as the mean of IrSPI-induced percentage inhibition after subtraction of the refolding buffer effect and comparison with the control. Double asterisks (**) indicate significant results as determined via ANCOVA analysis (P<0.001)

### IrSPI expression and localisation

To explore IrSPI function, we analysed its expression by quantitative RT-PCR in all tick stages from our pathogen-free *I. ricinus* colony. IrSPI mRNA was not detected in eggs, unfed larvae, nymphs, adults of either sex, or in fed larvae at days 1 and 3 of engorgement. However, comparison of transcript abundance in unfed, 3-day fed, and 5-day fully engorged nymphs revealed significant upregulation of IrSPI mRNA following 5 days of feeding (Fig 3A). IrSPI mRNA expression was then evaluated in different tissues in a pool of 17 field-collected females pre-fed for 5 days on a rabbit that was shown to be infected with *Babesia venatorum* and *Rickettsia helvetica* by PCR as previously described [8]. Results showed that in adults, and at roughly half way through feeding, *IrSPI* is expressed in SG, guts, synganglion, ovaries and the rest of tick bodies (carcasses), with significantly higher expression in SG (Fig 3B). *IrSPI*-specific transcripts were then analysed in various organs of *B. henselae*-infected *I. ricinus.* Results confirmed over-expression in SG following infection, and also revealed upregulation in gut, and in ovaries to a lesser extent (Fig 3C). Such overexpression was not found following infection with *B. birtlesii,* a phylogenetically proximal bacterium.

**Fig 3.**
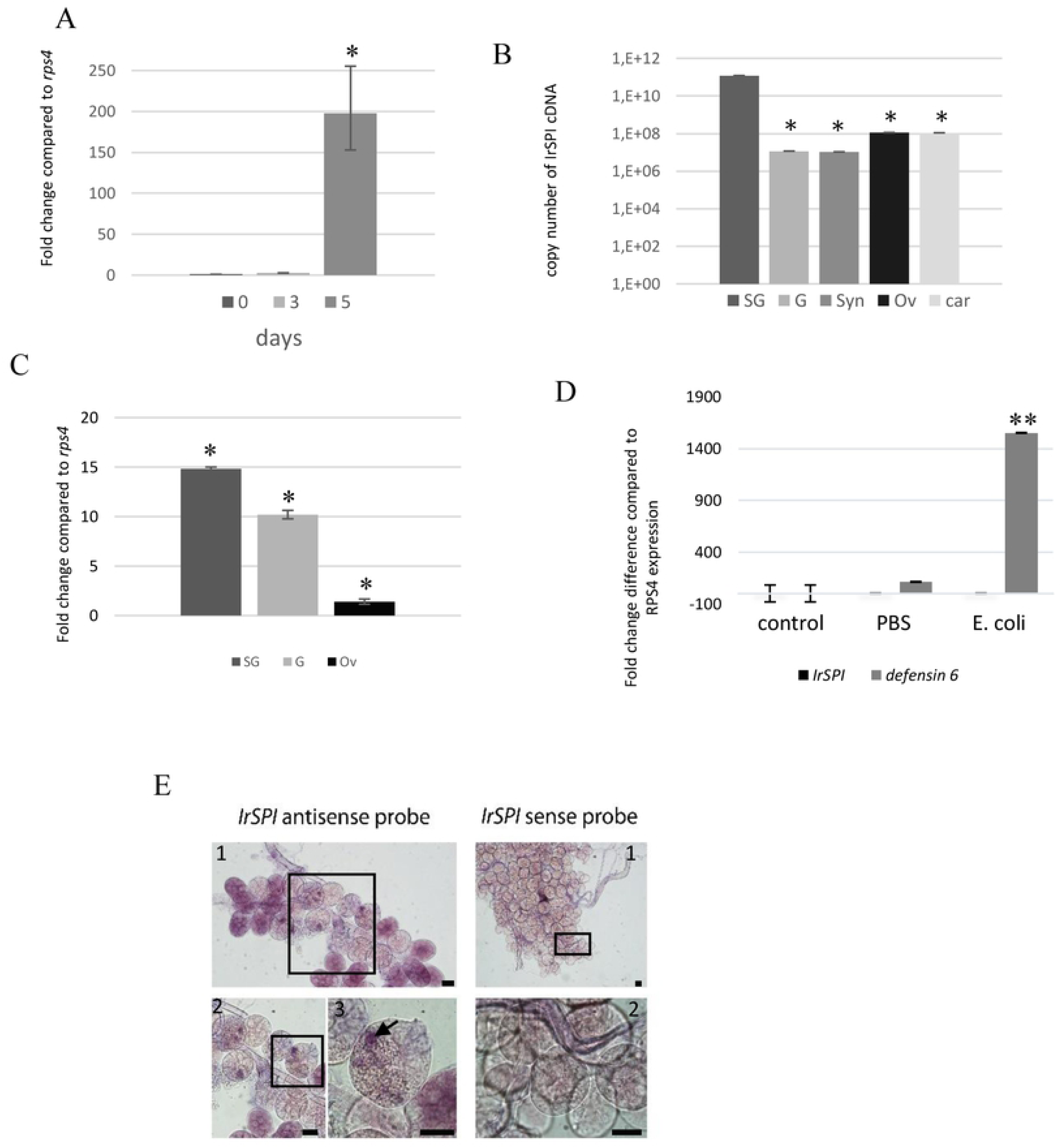
*IrSPI* transcript profiling and localisation. A) Relative *IrSPI* expression in pathogen-free nymphs fed on rabbits revealing the delayed gene expression at day 5. At each feeding time point (Days 0, 3, and 5), 20 pathogen-free nymphs were used, and *IrSPI* expression was assessed by quantitative RT-PCR relative to the *rps4* gene. B) Relative *IrSPI* expression in several organs of wild female ticks pre-fed for 5 days on rabbit. Pools of salivary glands (SG), gut (G), synganglion (Syn), ovaries (Ov), and the remaining body, i.e. the carcasses (car) from 17 females were used for each condition, and *IrSPI* expression was assessed by quantitative RT-PCR according to the copy number of *IrSPI* plasmid clones. C) Relative expression of *IrSPI* in pools of 13 salivary glands (SG), 14 guts (G), and 16 ovaries (Ov) from *B. henselae*-infected females pre-fed for 4 days assessed by quantitative RT-PCR, compared with uninfected samples, and expressed as a fold change with respect to the *rps4* gene. D) Relative *IrSPI* and *defensin6* expression after *E. coli* injection. Three pathogen-free females per group either received no injection (Control), 250 nl of PBS (PBS), or 250 nl of 10^7^ bacterial CFU (*E. coli*). After 24h quantitative RT-PCR was used to assess the mean expression of *IrSPI* and *defensin 6* normalised to *rps4*. E) *In situ* hybridization of *IrSPI* mRNA in salivary glands of pathogen-free females pre-fed for 8 days on membrane. A specific *IrSPI* anti-sense probe was incubated on salivary glands and visualised through addition of NBT/BCIP. Arrows indicate the presence of *IrSPI* transcripts in cells containing secretory vesicles in type II acini. An *IrSPI* sense probe was used as a negative control. Fields 1 to 3 are successive zooms. The scale bars are 60 µm. One asterisk (*) indicate significant (P<0.05) and two asterisks (**) highly significant (P<0.01) differences as determined by Student’s t-test.

*IrSPI* expression was also analysed after infecting ticks with a non-transmitted bacterium, *Escherichia coli,* to evaluate the protein’s potential implication in tick innate immune responses. The efficiency of this response was validated in parallel by evaluating expression of *defensin6*, which is known to be upregulated in ticks following *E. coli* infection [9]. While we confirmed that *defensin6* was upregulated following infection, expression of *IrSPI* remained unchanged (Fig 3D).

To document *IrSPI* expression in the salivary glands and to determine whether the gene product is present in tick saliva injected into the vertebrate host, *IrSPI* mRNAs were analysed by *in situ* hybridization in salivary glands of pathogen-free, partially-fed females. The *IrSPI* transcript was identified in the secretion vesicles of type II acini, which are responsible for saliva secretion in hard ticks [10] (Fig 3E). The presence of IrSPI protein was then evaluated in tick saliva and in salivary gland extracts obtained from a pool of 17 *I. ricinus* females collected from the field, then pre-fed on a rabbit for five days. SDS-PAGE analysis revealed major differences in the protein composition of saliva and salivary gland extracts, with higher total protein observed in salivary glands (Fig 4A). The presence of IrSPI was confirmed by western blot analysis using antiserum from mice immunised with recombinant IrSPI (Fig 4B), as well as serum collected from an experimentally tick-infested rabbit (Fig 4C). Native IrSPI was only faintly detected by the mouse antiserum in tick salivary glands, and not at all in saliva (Fig 4B). However, the recombinant counterpart was detected by the serum from a tick-infested rabbit, demonstrating that IrSPI generates an antibody response in rabbits, and is indeed injected into the host by ticks *via* the saliva during blood feeding (Fig 4C). These results suggest possible differences regarding immunogenicity/conformation and/or protein quantity between the recombinant and native proteins, since mouse antiserum recognised lower levels of native versus recombinant IrSPI.

**Fig 4.**
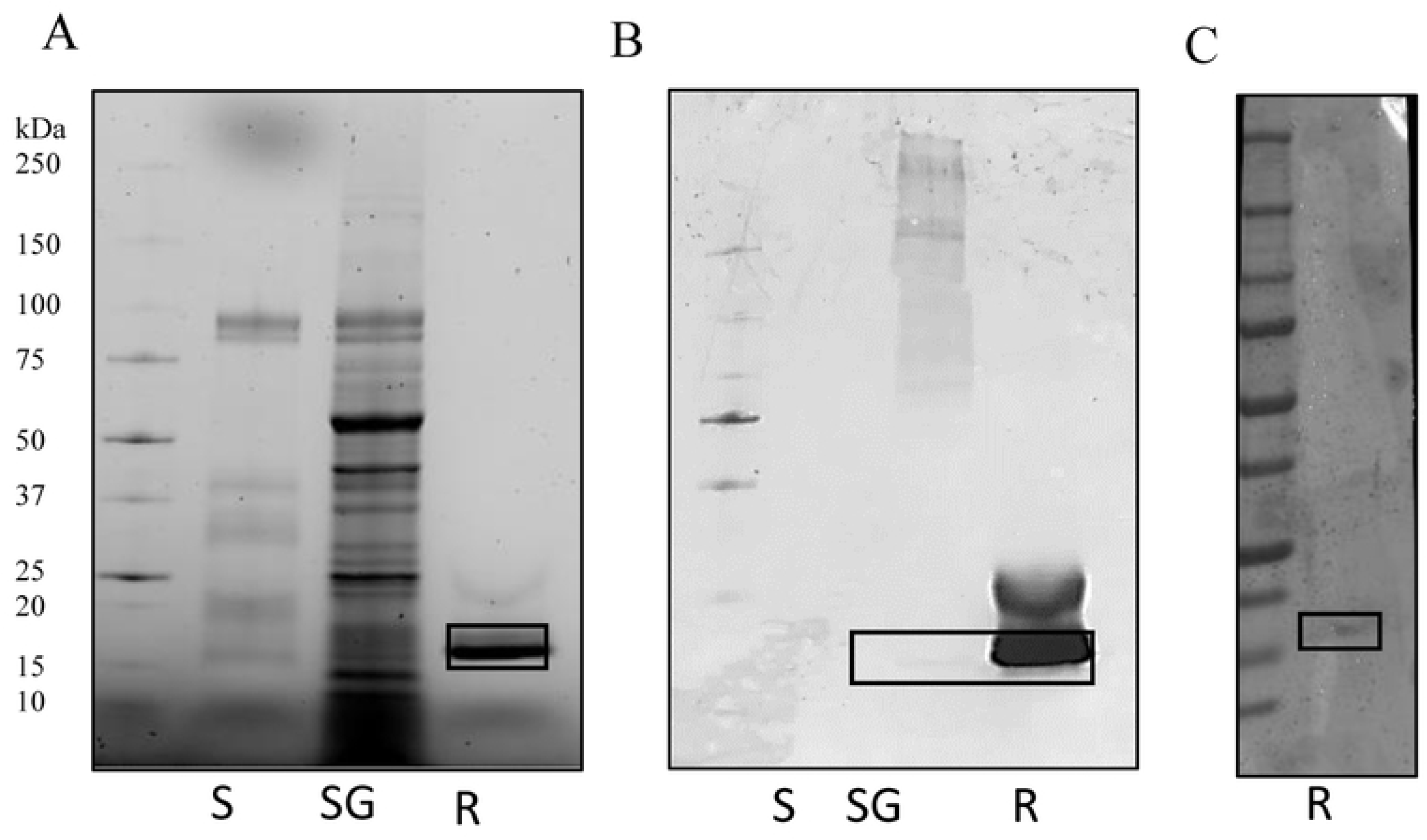
Detection of endogenous IrSPI protein expression. Tick saliva (S) and salivary gland extracts (SG) were obtained from a pool of 17 *I. ricinus* females from the field after a five-day pre-feeding step on rabbits. To each lane, samples of 10 µl of saliva (S), 10 µl of salivary gland extract (SG), and 1.4 µg of recombinant IrSPI (R) were loaded. A) SDS-PAGE electrophoresis and UV visualisation. B) Western blot analysis using an antiserum from an IrSPI-immunised mouse. C) Western blot analysis using an antiserum from a tick-infested rabbit. Bands corresponding to IrSPI are framed with boxes.

### Anticoagulant activity

Thrombin generation and clot waveform assays were used to test whether IrSPI displays anti-coagulant activity. Thrombin formation was triggered by adding calcium to a premix of plasma, phospholipid vesicles, and tissue factor, in the presence or absence of varying amounts of ‘refolded’ IrSPI. The kinetic analysis suggested that IrSPI had little or no impact on clot formation (Fig 5). At concentrations of up to 0.8 µM IrSPI, clot formation lag-time and amounts of thrombin formed were comparable between the positive control (peak at 371 s) and samples containing IrSPI (peak at 371 s), or refolding buffer alone (peak at 357 s, Fig 5A). IrSPI also had no significant impact on fibrin formation, with respective clotting-times of 105 s for controls (with or without refolding buffer) versus 119 s for samples containing IrSPI (Fig 5B). IrSPI’s clot lysing ability was also investigated to evaluate its effect on clot stability. Fibrin formation was triggered as above—with a mixture of phospholipid vesicles, tissue factor, and calcium but in the presence of tissue plasminogen activator (tPA) to induce fibrin clot lysis. Again, there was no significant difference between half-lysis times using refolding buffer with or without IrSPI (512 versus 528 s, respectively; Fig 5C).

**Fig 5.**
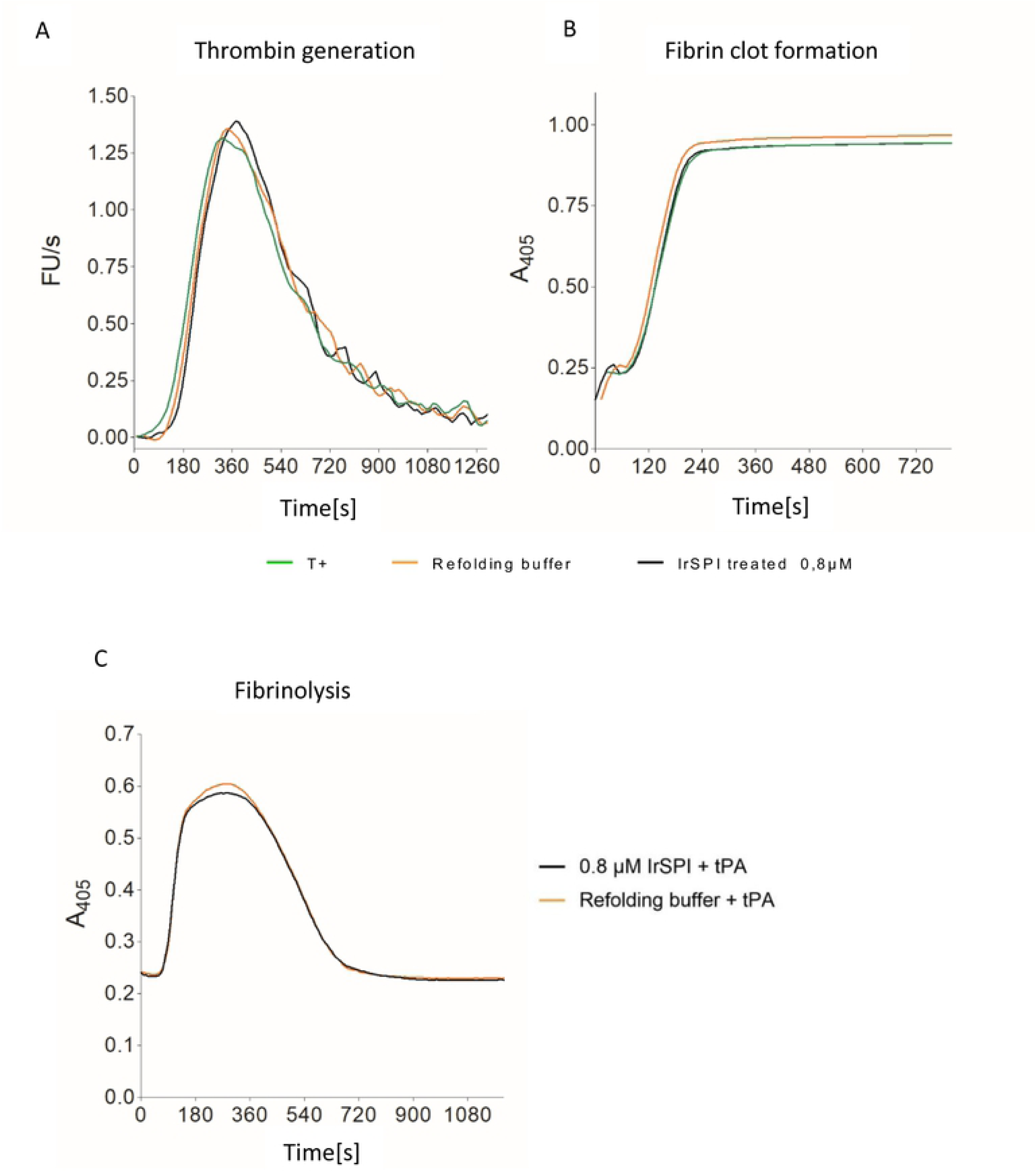
Effect of IrSPI on vertebrate host coagulation. A) Thrombin generation (in fluorescence unit/s) triggered by adding 10 mM CaCl2 to platelet poor plasma containing tissue factor and phospholipid (PPP-reagent). Curves in black were obtained in the presence of 0.8 µM IrSPI in refolding buffer, curves in orange in refolding buffer only, and curves in green in 0.15M NaCl. B) Simultaneous recording of the fibrin clot formation detected through the increase of turbidity at 405 nm; legend as in A). C) Fibrin clot formation was triggered as in A and B except that 6 nM tPA was added prior to clot formation. Half-lysis times (defined as the time to halve maximum turbidity) were not significantly different in the presence (curve in black) or absence (curve in orange) IrSPI. Results presented are mean of two independent experiments.

### Apoptosis and Angiogenesis

As IrSPI is known to be a component of tick saliva, which precludes tissue remodelling and angiogenesis, we next hypothesised that it could modulate formation of new vessels by promoting apoptosis and/or by inhibiting microvascular endothelial cell proliferation. The ability of refolded or misfolded recombinant IrSPI to induce apoptosis in comparison with cognate media controls, was first evaluated in HSkMEC endothelial cells by flow cytometry cell viability and apoptosis markers, i.e., cells labelled with Annexin V-FITC but not propidium iodide corresponded to apoptotic cells (Fig 6). Adding 1 µM IrSPI (refolded or not) to culture medium had only a weak effect (11.9% and 9.2% apoptotic cells versus 10.5 % with culture medium alone); and the greatest effect was actually obtained by the addition of refolding buffer without IrSPI (25.5%).

**Fig 6.**
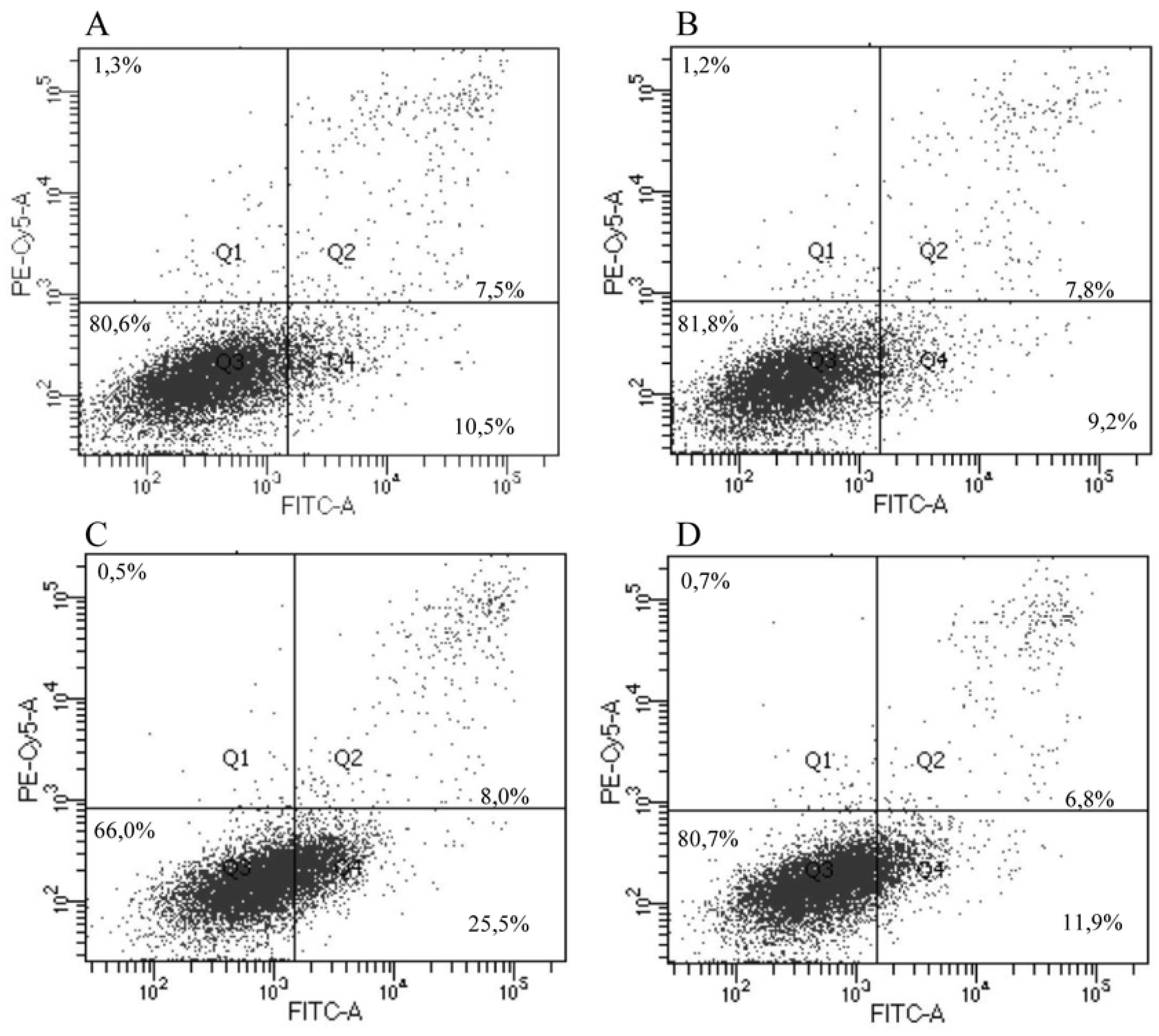
Impact of IrSPI on endothelial cell apoptosis. Viability and apoptosis were monitored by flow cytometry in HSkMEC endothelial cells after labelling with propidium iodide and Annexin V-FITC, respectively. Cells were incubated for 40 h in the presence of A) culture medium, B) 1 µM of IrSPI without refolding, C) refolding medium, or D) 1 µM of refolded IrSPI. The percentage of apoptotic cells was determined as the number of cells labelled with Annexin V but not propidium iodide (Q4) out of 10000 cells analysed per condition.

Two endothelial cell lines were used to investigate the impact of IrSPI on angiogenesis: HSkMEC and HEPC-CB1. We first evaluated IrSPI toxicity (with dilutions between 0.0625 and 2 µM, refolded or not) as well as culture- or refolding-media toxicity on HSkMEC cells. Concentrations of refolded IrSPI (0.5 µM and 1 µM) that did not induce significant toxicity compared to control (mortality of 18% and 29%, respectively), were used to evaluate IrSPI’s impact on angiogenesis. Compared with culture medium, non-refolded IrSPI at either concentration had no effect on the number of junctions, master junctions, segments, master segments, meshes, or total segment lengths in either of the two endothelial cell lines (Fig 7). In both cell lines, both concentrations of refolded IrSPI reduced several angiogenesis parameters, but this effect was mainly due to the refolding buffer and not IrSPI itself (Fig 7).

**Fig 7.**
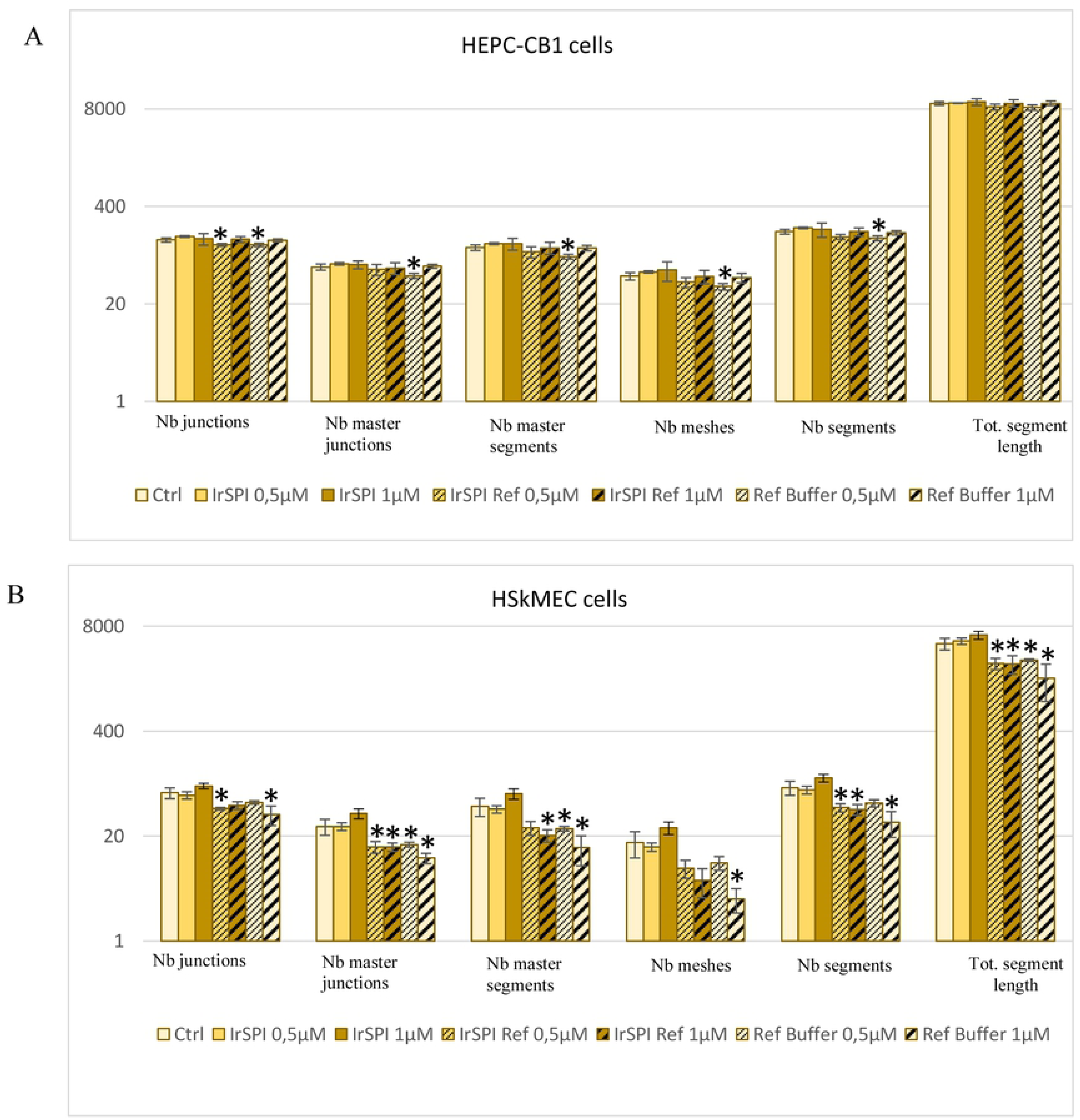
Impact of IrSPI on angiogenesis. A) HEPC-CB1 and B) HSkMEC endothelial cells were cultivated with IrSPI prior to (IrSPI), or after a refolding step (IrSPI Ref) for 9.5 h, and the number of junctions, master junctions, segments, master segments, meshes, and total segment lengths were recorded *via* video-microscopy every 30 min and analysed with ImageJ software. Results are expressed as means of triplicates. Controls correspond to culture medium (ctrl) and refolding buffer (Ref buffer). Asterisks (*) indicate significant differences as determined using the Student’s t test (P<0.05).

### Splenocyte proliferation assay

Next, we evaluated whether IrSPI might play a role in host immunomodulation during tick feeding. Splenocytes isolated from three OF1 mice were cultured in the presence or absence of the mitogen concanavalin A (ConA) for three days. The impact of non-refolded IrSPI on the proliferation of three lymphocyte subsets (CD4^+^ and CD8^+^ T cells, B cells) was then evaluated by CFSE dye dilution assay *via* flow cytometry. In the absence of IrSPI, mitogen-stimulated T but not B cells displayed robust proliferation, though with some inter-individual variability probably attributable to the use of outbred mice (Fig 8). In the presence of IrSPI, the proliferation of the global T cell population diminished, as evidenced by increased frequencies of undivided cells and decreased frequencies of cells that had undergone multiple divisions. Much of the observed reduction in proliferative capacity could be attributed to the CD4^+^ T cell subset; indeed, in the presence of IrSPI, the diminution in the frequency of cells having undergone one or more divisions was statistically significant in CD4^+^ (18%) but not in CD8^+^ (3.6%) T cells. Thus, IrSPI diminished the capacity of T cells, and especially that of the CD4^+^ subset, to respond to mitogenic stimulus, thus suggesting a role for IrSPI in immune modulation.

**Fig 8.**
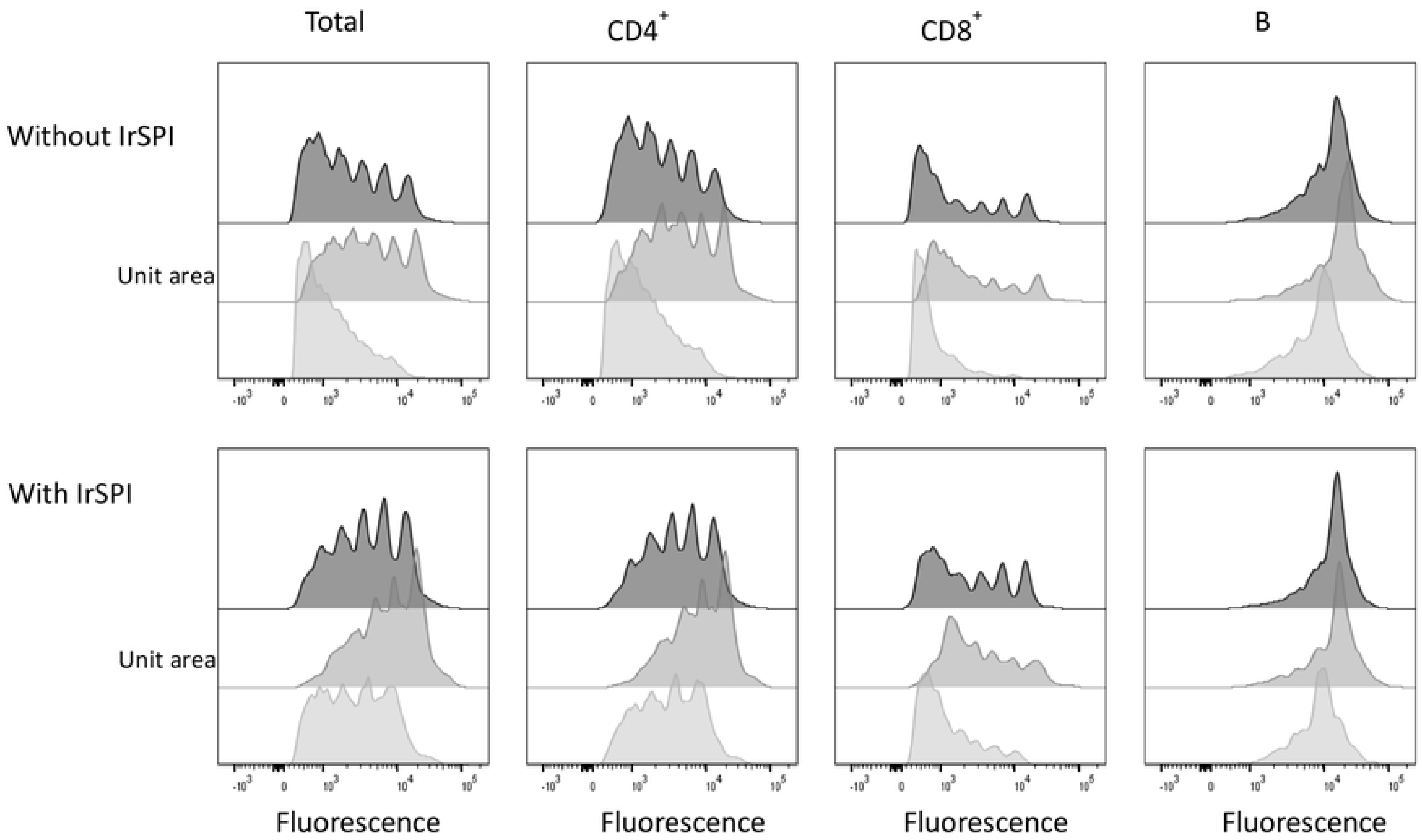
Impact of IrSPI on murine splenocyte proliferation. The impact of IrSPI on proliferation of T and B cells subsets was evaluated in splenocytes of OFI mice following mitogenic stimulation by Concanavalin A. Cell proliferation was measured by CFSE dye dilution and flow cytometry. Splenocytes from each of 3 mice were separately evaluated in triplicate for each condition. Results are expressed for each condition as the mean of triplicate wells for each condition. Successive peaks, displaying diminished CFSE labelling, reveal successive cell divisions.

### Cytokine expression in response to IrSPI

In order to gain insight into how IrSPI might influence immune cells, cytokines and chemokines secreted into the supernatants of mitogen-stimulated or unstimulated splenocytes were compared using a multiplex bead-based assay. The 26-plex panel included 17 cytokines representing multiple T helper subsets (Th1, Th2, Th9, Th17, Th22, and Treg) and 9 chemokines. In the absence of mitogen stimulation but in the presence of IrSPI, the secretion of several cytokines and chemokines was significantly diminished (P<0.01), in particular IP10 (−32.8%), MCP3 (−72.3%), MIP1β (−26%), and RANTES (−15.6%) (Table 1 and S1). IrSPI also significantly decreased several chemokines in mitogen-stimulated splenocytes including IP10 (−46.5%), MIP1β (−17.9%), RANTES (−37.9%), GM-CSF (−71.9%), and Eotaxin (−21.3%) (P<0.01). Moreover, expression of several T helper cytokines was also significantly reduced, namely, IFN-γ (−50.3%), IL-1β (−45%), IL-18 (−46%), IL-13 (−75.1%), IL-6 (−54.8%), TNF-α (−46,1%), and IL-9 (−48.8%) (P<0.01), Tables 1 and S2). In contrast, a single cytokine, IL-2, was significantly upregulated (+49.2%) following IrSPI exposure.

**Table 1:**
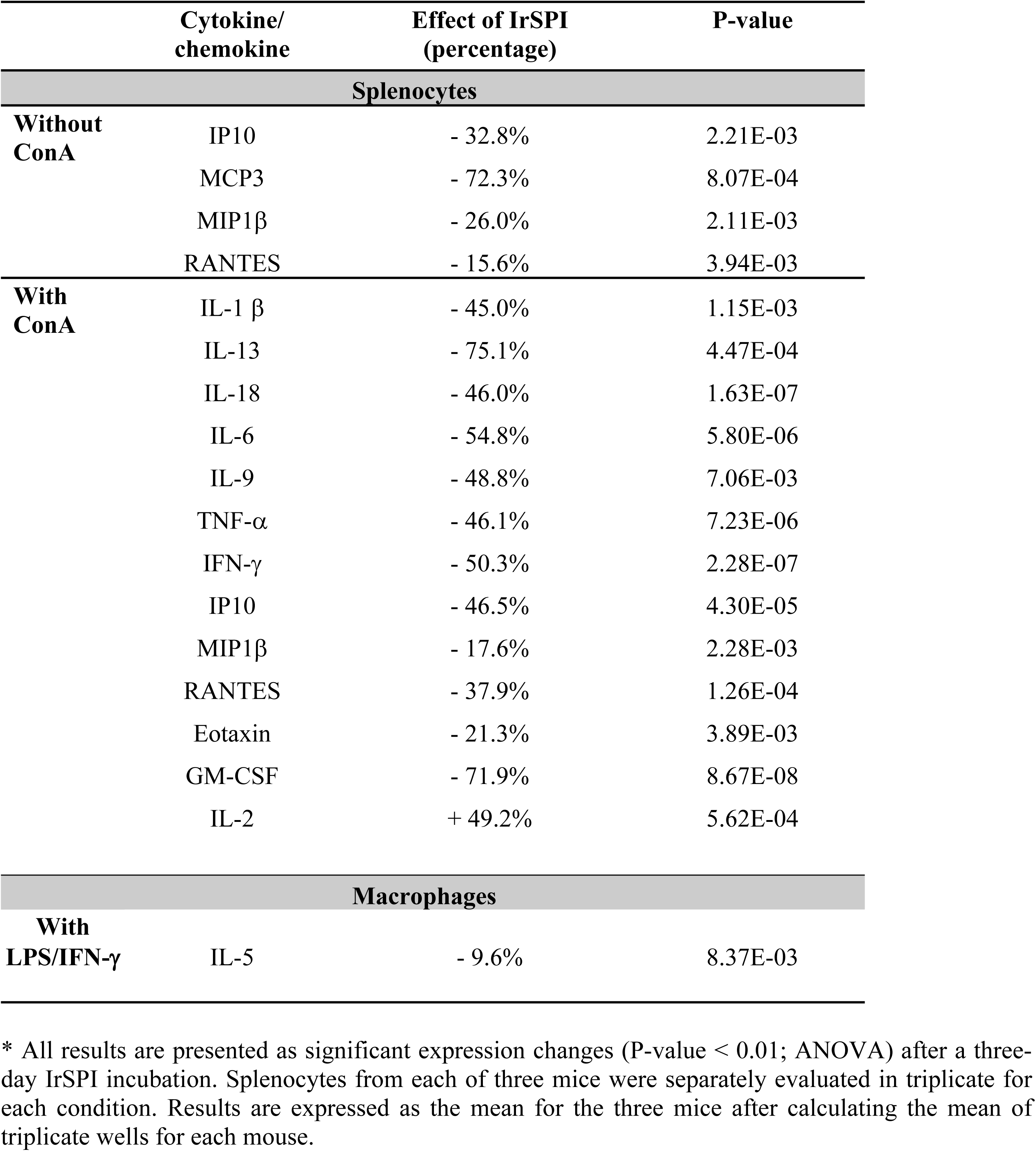
Impact of IrSPI on murine splenocyte cytokine/chemokine production with or without mitogen stimulation by concanavalin A (ConA).

To address the impact of IrSPI on sentinel immune cells, we also compared T helper cytokines and chemokines in the supernatants of IFN-γ- and LPS-activated murine macrophages grown in the presence or absence of IrSPI. All tested T helper cytokines and chemokines were diminished, but only the decrease in IL-5 was statistically significant (−9.6%; P<0.01; Tables 1 and S3).

Taken together, our data suggest that, apart from IL-2, IrSPI repressed splenocyte- and macrophage-activated secretion of multiple T helper cytokines and chemokines, consistent with a role for IrSPI as a modulator of the host immune response against tick bites.

## Discussion

Further knowledge is continually accumulating regarding the capacity of hard and soft tick saliva to counteract native host immune responses to enable successful blood meal completion [11–13]. Relatively few salivary compounds, however, have undergone thorough molecular characterisation, or have been identified as the active factors responsible for this process (recently reviewed in [3]). We previously identified a tick serine protease inhibitor—IrSPI—potentially present in *I. ricinus* saliva and likely implicated in tick feeding and in *I. ricinus* salivary gland infection by *Bartonella* [5]. The presence of genes homologous/paralogous to *IrSPI* in both *I. ricinus* and *I. scapularis* ticks suggests that it belongs to a family of SPI proteins that have important physiological functions, with highly conserved regulatory roles. Here we also provide evidence that IrSPI is a Kunitz SPI that is induced during blood feeding and is present in tick saliva. Moreover, we show that IrSPI inhibits elastase and modulates host immune responses. It is likely that such immunomodulation substantially contributes to successful feeding and engorgement of *I. ricinus* ticks, and thereby facilitates TBP transmission.

Production of functionally active recombinant protein is known to be difficult. As an example, the *Dermacentor andersoni* p36 immunosuppressive salivary gland protein was only active when the recombinant protein was expressed in insect cells, but not in bacteria [14], suggesting that functional activity might depend upon proper post-translational modifications such as glycosylation. Similarly, Konnai and co-workers have tentatively attributed the inability of *Ixodes persulcatus* lipocalin to bind histamine to expression of recombinant protein in *E. coli* [15]. In the present study, we expressed IrSPI in *Drosophila* cells, and showed that the protein was functional, e.g., it exerted immunomodulatory activity. However, circular dichroism analysis of recombinant IrSPI suggested that it lacked important Kunitz secondary structural features. Nonetheless, when subjected to a refolding step, the recombinant protein displayed inhibitory activity against elastase.

In accordance with its primary structure, IrSPI belongs to the Kunitz SPI superfamily, whose members competitively inhibit serine protease activity in a reversible lock-and-key fashion. In ticks, certain members of this superfamily are known to subvert host defence mechanisms during feeding [7]. Invertebrate Kunitz inhibitors possess diverse inhibitory effects on several serine proteases depending on their amino acid sequence, especially the P1 residue of the reactive site loop. Positively charged Lys or Arg, as P1 residues are associated with inhibition of trypsin-like serine proteases; large hydrophobic residues, such as Phe, Tyr, or Leu, with chymotrypsin-like serine proteases; and small hydrophobic residues, such as Ala or Val, with elastase-like serine proteases [16]. We showed that IrSPI, harboring Ala as P1 residue does indeed inhibit elastase. IrSPI also inhibited chymotrypsin activity, albeit weakly, in keeping with the partial fit of an Ala-bearing SPI within the protease active site, but less effectively than an SPI bearing large hydrophobic amino acids at this P1 position. Elastase, a serine protease synthesised and released by neutrophil cells following trauma, is responsible for tissue inflammation and remodelling [17]; chymotrypsin also participates in this latter process [18]. Through its inhibitory activity, IrSPI might thus reduce inflammation and inhibit tissue repair processes following tick bite, as has been suggested for other Kunitz SPIs, such as *Ixodes scapularis* tryptogalinin which also inhibits elastase, among other proteases [19].

We showed that *IrSPI* mRNA was expressed in diverse organs from pooled ticks, including SG, gut, ovaries, and synganglion, as well as the carcasses of wild pre-fed ticks that harboured—for at least one of the pooled ticks—two TBP. This broad tissue distribution suggests that IrSPI might be implicated in multiple aspects of tick biology. *IrSPI* expression in pools containing infected ticks is compatible with its induction by TBP, as we previously demonstrated for *B. henselae* [5]. In fact, TBP infection has already been shown to modulate expression of many tick genes, and a number of these—including some encoding Kunitz proteins—have been identified as implicated in TBP transmission [20]. Such molecular interactions reflect the vector competence of an arthropod [4], and may be TBP-specific, as shown for IrSPI regarding *B. henselae* and *B. birtlesii*. Previous RNAi experiments have also shown that IrSPI is likely involved in bacterial adhesion, invasion, and/or multiplication within tick salivary glands, and also potentially in tick defence mechanisms, as it may act against other bacteria in competition with *B. henselae* [5]. This latter hypothesis was thus evaluated using *E. coli,* but *IrSPI* upregulation was not observed following tick infection with this non-transmitted bacterium.

We next demonstrated that *IrSPI* expression is induced in tick salivary glands by the blood feeding process, both in nymphs at the end of engorgement (day 5) and in females in the middle of the feeding process, and in the latter context at day 5 on rabbits, and at days 4 and 8 on membrane [5]. Similar blood-meal induction has been also reported for the *Irsgmg-150466* transcript, which likely corresponds to the *IrSPI* gene as it has 100% identity at the protein level, but without a signal peptide sequence. *Irsgmg-150466* expression peaked 12 h after attachment, but then decreased after 24 h and 12 h in *I. ricinus* female and nymph salivary glands, respectively [21]. Chmelar and co-workers reported the expression of a homologous gene (contig 240) in *I. ricinus* female salivary glands 24 h after tick attachment [22]. Dynamic expression of IrSPI paralogs was also recently reported by Dai *et al*. in *I. scapularis* [23], lending support to variable BPTI/Kunitz gene expression by both *I. ricinus* and *I. scapularis* during blood meal ingestion, with time-dependent up- or down-regulation. Nevertheless, it should be noted that in these studies, expression level is only based on mRNA detection, which may differ from protein expression, as variable transcript expression does not necessarily translate into variation in expression of the corresponding protein.

The presence of a signal peptide combined with salivary gland transcript expression indicated that IrSPI was likely to be a secreted protein present in tick saliva. We have shown that *IrSPI* mRNA is expressed in type 2 acini, which are responsible for saliva secretion, and, using anti-IrSPI antibodies raised in tick-infested rabbits, that native IrSPI protein is present in tick saliva and injected into the host during feeding. This demonstration that IrSPI takes part in the molecular dialogue between the tick and its vertebrate host, and strongly argues in favour of a role for IrSPI in the tick feeding process.

Host skin is the first barrier that ticks need to overcome for successful feeding. Upon tissue damage, host skin cellular components—including epithelial cells, fibroblasts, and endothelial cells— generate biomolecules that facilitate tissue remodelling and wound healing, such as angiogenesis. In order to remain attached to their host and capable of obtaining blood over several days, these slow-feeding hematophagous arthropods must inhibit these processes. The first study demonstrating that tick saliva inhibits angiogenesis was published in 2005 [12]. Since then, a number of tick salivary molecules have been reported to modulate angiogenesis and wound healing. These include Kunitz SPI from *Haemaphysalis longicornis* (Haemangin [24]), *R. microplus* (BmTI-A [25] and BmCI [26]), and *Amblyomma cajennense* (Amblyomin-X [27]). Nevertheless, although IrSPI, like BmTI-A and BmCI, acts as an elastase inhibitor, it does not seem to exert direct activity on apoptosis of HSkMEC cells, or angiogenesis in HSkMEC and HEPC-CB1 cells. It should be noted, however, that this *in vitro* observation may not hold true for the whole organism, where suppression of angiogenesis-related proinflammatory cytokines by IrSPI, as observed in both splenocytes and macrophage cells, might exert an indirect effect.

To initiate blood feeding, the tick must lacerate the host tissue, which exposes to blood large quantities of tissue factors triggering activation of the blood coagulation cascade that induces fibrin clot formation and its subsequent lysis. Blood coagulation is thus rapidly triggered following tick bite-induced blood vessel damage. Several tick factors counteract vertebrate blood haemostasis (reviewed in [3]), the majority belonging to the SPI family. In fact, most of the tick SPI injected into the host during blood feeding facilitate completion of the long blood meal by targeting one or several serine protease(s) involved in blood coagulation pathways, thereby preventing fibrin clot formation and maintaining blood in a fluid state at the tick bite site [7]. IrSPI, however, did not appear to modulate blood coagulation, as no effect was observed on thrombin generation or fibrin formation. In addition, albeit several tick SPI accelerate lysis of fibrin clot [7], IrSPI failed to demonstrate such activity. Nevertheless, result was somehow expected given that the P1 residue of IrSPI is alanine suggesting elastase-like targets rather than blood coagulation proteases which are all trypsin-like enzymes [28]. Further investigations would be required to determine whether IrSPI regulate other blood coagulation processes, such as platelet aggregation.

To avoid host rejection, ticks have developed a complex armament to circumvent both innate and adaptive immune responses as well as complement activation [11]. Relatively few tick salivary factors responsible for this immunomodulation have been identified, (reviewed in [3]). These factors have been reported to play a role in the activation and/or recruitment of multiple resident and infiltrating cell types at the tick bite site, including macrophages, mast cells, dendritic cells, endothelial cells, keratinocytes, and lymphocytes, as well as in the polarisation of an initial Th1 helper response towards a Th2 orientation, considered to be more favourable to tick blood feeding [11]. Here we showed that IrSPI inhibits mitogen-induced proliferation of murine CD4^+^ T cells, as has been reported for saliva of not only *I. ricinus* [29], but also several other tick species including *D. andersoni* [30], *A. variegatum* [31], and *I. dammini* [32]. It is noteworthy that in our study, IrSPI inhibited secretion of IL-9, which promotes the proliferation of CD4^+^ rather than CD8^+^ T cells [33]. Until now, a small number of salivary proteins with immunosuppressive capacity have been characterised, including Salp15 from *I. scapularis* [34], the most highly-investigated protein, P36 from *D. andersoni* [14], and Iris from *I. ricinus* [35]. It should be noted that in our study, the impact of IrSPI was assessed on mouse splenocytes, which comprise, in addition to the high proportion of T cells, multiple other immune cell types such as B cells, macrophages, dendritic cells, eosinophils, basophils, natural killer cells and neutrophils. Thus, indirect actions of IrSPI on T cells *via* other cells should not be excluded, as has already been suggested for dendritic cells [36]. Indeed, we observed that IrSPI decreased splenocyte expression of multiple chemokines and cytokines that directly or indirectly stimulate T cell proliferation, including MIP-1β, RANTES, and IL-9 [37, 38]. Furthermore, we showed that IrSPI also repressed the expression of IL-13, which is mainly produced by Th2 lymphocytes, and which may regulate proliferation and differentiation of DCs and B cells [39].

At the tick bite site, and in response to tissue damage, numerous cell types (including fibroblasts, endothelial cells, macrophages, DCs, T cells, neutrophils, eosinophils, and resident tissue cells) are able to secrete immunological factors, thereby promoting inflammation, tissue remodelling, wound healing, angiogenesis, and pathogen clearance. To repress immune cell activation and proliferation that, if unchecked, would compromise tick feeding, tick saliva, as a rich source of immune modulators (reviewed in [3]) acting in concert to achieve a stronger effect, hijacks cellular immune responses and notably secretion of cytokines and chemokines, thus affecting the previously mentioned processes. Our results showed that for *I. ricinus,* IrSPI, as part of this salivary cocktail, represents one of the molecules that is likely to contribute to the modification of these molecule profile, and suggest a larger role of IrSPI along tick feeding *in vivo* due to their numerous roles in mammalian physiology.

The capacity of tick saliva to polarise immune responses towards a Th2 rather than Th1 profile has been reported for several tick species including *I. ricinus* [36, 40, 41]. Our results show that by repressing the production of IP10, MCP 3, MIP-1β, and RANTES from a large variety of cells (present in splenocytes), IrSPI may actively contribute to Th2 immune polarisation by restraining Th1 cytokines. Another elastase inhibitor from *I. ricinus*, Iris, has been shown to promote Th2 polarisation by enhancing Th2 cytokine expression at the tick bite site by increasing IL-10 secretion and repressing IL-6 and IFNγ [35]. It has been reported that polarisation of the immune response toward the Th2 subset is associated with tolerance to tick bites in mice [42], in addition to creating favourable conditions for the tick-borne bacteria *Borrelia burgdorferi* [43].

This Th2 orientation is expected to promote an anti-inflammatory context, and indeed inhibition of pro-inflammatory molecules by total salivary gland extracts of *I. ricinus* has already been reported [40]. Among IrSPI down-regulated molecules, MIP-1β, GM-CSF, and RANTES—strongly associated with secretion of IFN-γ—are considered to be proinflammatory chemokines typically expressed in response to infection, promoting recruitment of effector cells to the site of pathogen entry [44]. Our data also showed that IrSPI decreased the expression of IFNγ, as has been reported for salivary gland extracts of *I. ricinus* [45]. IFNγ is principally produced by cytotoxic and Th1 cells, and constitutes one of the major regulators of both innate and adaptive immunity, as well as inflammation. Local tissue inflammation and allergic reactions at the tick bite site are also controlled by IrSPI through the inhibition of eotaxin-1, IL-18, or IL-9. Inhibition of this last cytokine has also been reported for the salivary cysteine protease inhibitor sialostatin L from *I. scapularis*, thus preventing the development of experimental asthma [46].

In addition, and like the dendritic cell modulator Japanin from *Rhipicephalus appendiculatus* [47], IrSPI also repressed the expression of TNFα, which is predominantly produced by both activated macrophages and T lymphocytes in inflammatory and infectious conditions. Although unable to drive the differentiation of naïve CD4^+^ cells towards a Th2 phenotype in a direct manner, IL-13, also repressed by IrSPI, may affect T cell function indirectly *via* downregulation of proinflammatory Th1 cytokines, including IL-12, which have been shown to be also repressed by OmC2 from the soft tick *Ornithodoros moubata* [48].

Like Japanin [47], IrSPI decreased IL-1β. In addition to triggering the release of other pro-inflammatory cytokines, this cytokine—which is secreted following injury—activates neutrophil and macrophage phagocytosis and the release of toxic oxygen and nitrogen radicals. As serine proteases from neutrophils and macrophages, such as elastase, have been reported to process pro-IL-1β into active fragments [49], we thus cannot exclude a role for IrSPI as an indirect inhibitor of IL-1β secretion through inhibition of these molecules. As this cytokine was among the most highly downregulated in the presence of IrSPI, and given the importance of IL-1β in both immune cell recruitment and nociception, its suppression by IrSPI may well participate in avoidance of tick rejection by the host. Besides IL-1β, pain sensation is reinforced by release of pro nociceptive chemokines and cytokines, including TNFα, IFNγ, IL-6, IL-18, MCP3, RANTES, and MIP-1β, at the tick bite site [50], and these are also repressed by IrSPI in cultured splenocytes. Immune cell recruitment might also be impacted by IrSPI through the modulation of MIP-1β, RANTES, GM-CSF, and IP10 or IL-6, triggering the expression of T cell attractant chemokines and eotaxin-1, TNF-α, or IL-13 which regulate cell adhesion proteins [51].

As ticks must maintain blood flow at the bite site in order to feed successfully, inhibition of several cytokines/chemokines by IrSPI might antagonise host homeostasis and angiogenesis. Affected cytokines include RANTES, which modulates neovascularization, endothelial cell migration and spreading [52], IL-6, which induces VEGF [53], TNF which stimulates *COX 2* [54], and eotaxin-1 which affects endothelial cell chemotaxis and *in vivo* blood vessel formation [55].

The immunosuppressive effect of IrSPI on the host will naturally also influence transmission of tick-borne pathogens. Several of the cytokines/chemokines repressed by IrSPI, including RANTES, MIP-1β, IL-18 and IL-9, are also implicated in pathogen control and clearance mechanisms, including effector cell recruitment to the pathogen entry site ([44], [56]). While most proinflammatory cytokines and chemokines were significantly inhibited by IrSPI, a single cytokine, IL-2, was upregulated. This result was somewhat unexpected in the light of published data reporting decreased IL-2 expression in tick saliva [11]. Nevertheless, while IL-2 is often considered to be proinflammatory, it can also behave as an anti-inflammatory cytokine by promoting differentiation of Treg cells as well as their long-term survival, and suppressive activities [57], thereby decreasing local inflammation and promoting pathogen tolerance [58].

Together with the splenocyte results, the macrophage response to IrSPI confirms that this salivary molecule downregulates both proinflammatory and Th1-promoting cytokines. A similar result has been reported where TNF-α, IL-12, and IFN-γ were decreased in LPS-activated bovine macrophages exposed to *R. microplus* saliva [59]. While all tested cytokines decreased in this study, only IL-5 from mouse macrophages was significantly lowered. This cytokine is reported to have pro-inflammatory functions, including eosinophil activation, B cell growth, and the stimulation of antibody production. Its pro-inflammatory features are reinforced *via* its ability to increase the susceptibility of B and T cells to IL-2, and subsequently the production of cytotoxic T cells. Although this cytokine is predominantly expressed by CD4^+^ T cells, recent evidence suggests that macrophages may also produce IL-5 [60].

IrSPI is produced by the most common tick vector in Europe, and is a vital component of the tick salivary cocktail that enables the tick to overcome the host immune system’s defences to feed successfully and transmit pathogens. While further investigations are required to gain full understanding of the molecular processes that modulate the host immune response, this work represent a significant advance in the characterization of key immunomodulatory molecules in tick saliva. IrSPI also represents a potential target antigen for anti-tick vaccine development, as host immune responses elicited against such immunomodulatory tick molecules may diminish tick infestation and/or TBD transmission. Anti-tick vaccines hold the promise of affording broad protection against multiple TBD transmitted by the same vector. The utilisation of “exposed” salivary antigens such as IrSPI, rather than “concealed” tick antigens to which the host is never naturally exposed, may enable the host response to be naturally boosted upon exposure to ticks [61]. In addition, immunity against such proteins may block the feeding process prior to pathogen transmission, which typically occurs many hours or even days after tick attachment. Finally, and although we have reported an impact of *IrSPI* silencing on both tick feeding and bacteria transmission, IrSPI belongs to a protein family with extensive functional redundancy [62]. In consequence, the vaccinal potential of IrSPI would likely be evaluated by promoting immune responses against epitopes that are common to the IrSPI protein family. Moreover, the protection afforded by IrSPI should ideally evaluated in the context of multivalent anti-tick vaccines comprising a cocktail of salivary antigens involved in distinct mechanisms that counteract host defences by different mechanisms. Furthermore, due to its immunomodulatory activities, IrSPI may also represent an interesting candidate as a therapeutic molecule in pathologies linked to immune cell proliferation disorders.

## Material and methods

### Ethics

This study was carried out in strict accordance with good animal care practices recommended by the European guidelines. Animal experiments involving rabbits and mice used for tick feeding were approved by the local ethics committee for animal experimentation (ComEth Anses/ENVA/UPEC; Permit Number 2015091413472401). Rabbits used for feeding pathogen-free ticks have been put up for adoption *via* the White Rabbit Association, Paris, France.

### Ticks

This study used *I. ricinus* ticks either collected from the field, or from a pathogen-free colony. Questing ticks (nymphs and adults) were collected by flagging in the Sénart forest, located south-east of Paris (France), as previously described [8]. All pathogen-free *I. ricinus* ticks were derived from a laboratory colony reared at 22°C in 95% relative humidity, with 12 h light/dark cycles as previously described [63].

### Cloning and sequencing of *IrSPI* cDNA

Following NGS of the *I. ricinus* salivary gland transcriptome, we had previously generated a total of 24,539 isotigs [5]. Of these, the *IrSPI* coding sequence (GenBank KF531922.2) without the stop codon (282 bp) was amplified with primers ID23822F (5’-ACATCTACCGTTCAAGATGAAG-3’) and ID23822R (5’-TCTAAGTGCCTTGCAGTAGTC-3’), and then cloned into the pET28 vector following the manufacturer’s instructions (Novagen). After transformation of *E. coli* TOP10 bacteria (ThermoFisher Scientific) using plasmid and kanamycin selection, correct *IrSPI* sequence insertion was validated by PCR and sequencing.

### Recombinant IrSPI protein production

The open reading frame (ORF) of IrSPI lacking the 5’ sequence encoding its signal peptide (MKATLVAICFFAAVSYSMG) was synthetised (Eurofins Scientific) as a fusion protein with sequences encoding Twin-Strep-Tag (WSHPQFEKGGGSGGGSGGGSWSHPQFEK) and enterokinase (DDDDDK). The BglII and EcoRI restriction sites were used to insert the sequence into the pMT/BIP/V5-His plasmid (ThermoFisher). The production of recombinant IrSPI protein was performed by the Recombinant Proteins in Eukaryotic Cells Platform, Pasteur Institute, Paris, France. Briefly, Drosophila S2 cells (Invitrogen, Carlsbad, CA) adapted to serum-free Insect Xpress (Lonza) medium were co-transfected with pMT/BIP/IrSPI and the pCoPURO vector (Addgene) conferring resistance to puromycin, using the cellfectin II transfection reagent (Invitrogen). The upstream pMT/BiP/V5-His *Drosophila* BiP secretion signal enables the recombinant protein to enter the S2 cell secretory pathway for recovery in culture medium. IrSPI expression was driven by the metallothionein promoter after induction by 5 µM CdCl_2_. Clarified cell supernatants were concentrated 10-fold using Kicklab tangential flow filtration cassettes (GE Healthcare, cut-off 10 kDa) and adjusted to pH 8.0 in 10 mM Tris before purification in an AKTA Avant system by Steptrap-HP chromatography (GE Healthcare). Final purification was achieved by gel filtration using a Superdex 75 HiLoad 16/60 column (GE Healthcare) equilibrated in PBS-X (without Ca^2+^ and Mg^2+^). Protein quantity was estimated by peak integration.

### IrSPI refolding

Eight µg of IrSPI were refolded for 30 minutes at 25°C in a final volume of 100µl containing 10µg of the disulfide isomerase protein (Sigma-Aldrich) with reduced/oxidised glutathione (Sigma-Aldrich) (2mM/0.2mM) in borate buffer (100 mM Borate, 150 mM NaCl, pH 7.4).

### Circular dichroism

Circular dichroism experiments were performed in the far-UV region using an Aviv CD 215 model spectrometer equipped with a water-cooled Peltier unit. A 200µl volume of recombinant IrSPI at 0.075µg/µl was used to record spectra in a cell with a 1-mm path length in the far-UV range (between 190 to 270 nm) at 25°C. IrSPI and its cognate buffer were successively screened ten times to produce an averaged spectrum. The spectra were corrected using buffer baselines measured under the same conditions. Data were normalised to residue molar absorption measured in mdeg (M^−1^ cm^−1^) and expressed as delta epsilon (Δε) enabling the BestSel website algorithm to analyse for secondary structure predictions [64].

### Sedimentation velocity and analytical ultracentrifugation

The oligomerization state of IrSPI was determined in a sedimentation velocity experiment in a ProteomeLab XL-I analytical ultracentrifuge (Beckman-Coulter, USA). Briefly, after loading IrSPI at 0.075 mg.ml^-1^ in a 1.2 cm epoxy double-sector cell, the cell assembly was equilibrated for 1.5 hours at 20.0°C in a four-hole AN60-Ti rotor. The sample was spun at 42,000 r.p.m for 10 h and 400 scans for both Rayleigh interference and absorbance at 280 nm were recorded. Absorbance data were collected at a constant radial step size of 0.001 cm. Partial specific volume of 0.70 and the extinction coefficient ε at 280 nm of 14350 L.mol^-1^.cm^-1^ were theoretically calculated at 20 °C from the IrSPI amino acid sequence using Sednterp software (Spin Analytical, USA). PBS (-Ca^2+^ and -Mg^2+^) density ρ and viscosity η at 1.0053 g.mL^-1^ and 1.018 cP, respectively, were also determined with Sednterp at 20°C. The sedimentation profile of IrSPI was analysed in SEDFIT using the continuous size distribution c(s) model [65].

### MALDI-TOF/TOF analysis

Recombinant IrSPI protein purity and integrity were confirmed using a Bruker UltrafeXtreme MALDI-TOF/TOF instrument (Bruker-Daltonics, Germany). Briefly, 1 µL of protein at 0.075 mg/mL was deposited on a MTP 384 ground steel target plate with 1 μL of 2,5-dihydroxybenzoic acid (2,5-DHB) in 50% acetonitrile, 0.1 % trifluoroacetic acid as matrix solution. Data were acquired using Flexcontrol software (Bruker-Daltonics, Germany) and shots were recorded in positive ion linear mode. Mass spectra were externally calibrated in the m/z range of 5-20 kDa with a protein calibration standard (Bruker-Daltonics, Germany) and analysed with the Flexanalysis software (Bruker).

### Serine protease inhibition assays

Serine protease inhibition assays were performed at 37°C by using a thermostated Multiskan GO ELISA plate reader (Thermo Scientific). Serine proteases and their corresponding substrates (in brackets) were purchased from Sigma Aldrich and, serine proteases were used with the following concentrations: 24.6nM Trypsin-T1426 (N-Benzoyl-Phe-Val-Arg-*p*-nitroanilide hydrochloride), 46.5 nM Chymotrypsin-C3142 (N-Succinyl-Ala-Ala-Pro-Phe-p-nitroanilide), 21.6 nM Elastase-45125 (N-Succinyl-Ala-Ala-Ala-*p*-nitroanilide), 43nM Thrombin-T6884 (Sar-Pro-Arg-p-nitroanilide dihydrochloride), and 33nM Kallikrein-K3627 (N-Benzoyl-Pro-Phe-Arg-p-nitroanilide hydrochloride). Reactions were performed in 20 mM Tris-HCl, 150mM Nacl, 0.1% BSA, pH 7.4 buffer (except for thrombin inhibition assay performed at pH 6.5), with a final substrate concentration of 0.2 mM. All reactions were performed with both non-refolded and refolded recombinant IrSPI, the refolding buffer being used as a negative control for this latter condition. After initial incubation of serine protease with or without recombinant protein for 15 min at 37°C, the A_405_ was recorded for 30 minutes (every 10s) at the same temperature. Initial slopes of substrate hydrolysis together with their correlation coefficients were obtained by linear regression analysis using GraphPad Prism software (San Diego, CA, U.S.A.). Results were expressed as the percentage inhibition in the presence of IrSPI taking into account the effect of the refolding buffer alone on the enzyme activity.

### Ticks infected with tick-borne pathogens

*B. birtlesii*-infected nymphs pre-fed for three days on mice were obtained from moulted larvae infected on mice injected with an *in vitro B. birtlesii* (IBS325^T^) culture as previously described [66]. *B. henselae-*infected females pre-fed for four days were obtained with the previously described membrane feeding system and following the same protocol [5]*. B. henselae* and *B. birtlesii* infection in ticks was confirmed by RT-PCR using primers described by Reis and co-workers [66].

### Tick infection with *Escherichia coli*

*E. coli* (CIP7624 strain, Pasteur Institute, Paris) was cultivated at 37°C under atmospheric conditions for one night on Luria Bertani (LB)-agar plates and then for 24h in liquid LB medium. Bacterial concentration was then evaluated by spectrophotometry at 600 nm (OD at 1 represents 2.2 × 10^9^ CFU/ml). Three pathogen-free *I. ricinus* females were nano-injected with 250 µl (1 × 10^7^ CFU) of bacterial suspension into the body cavity using a glass microcapillary attached to a nanoinjector pump (Drummond) driven by a Micro 4 controller (World Precise Instruments). Three other females injected with 250 µl of PBS and three un-injected ticks were used as controls. *E. coli* infection was confirmed by qPCR with primers targeting the *WecA* gene [67].

### Tick saliva and tick organ collection

For saliva collection, 34 female *I. ricinus* from the field were pre-fed for five days on a rabbit as previously described [68]. After detachment, ticks were then physically immobilised on microscopy slides with the hypostome positioned in sterile 10 µl tips. After applying 5 µl 5% pilocarpine in methanol to the tick dorsum, ticks were positioned upside down to facilitate collection of saliva and placed in dark, humid chambers as described by Patton et al. [69]. After about 5h, a pool of approximately 50 µl of saliva was obtained (average of 1.5 µl per tick) and stored at −80°C until use.

After saliva collection, salivary gland extracts were prepared. Salivary glands were collected from 17 ticks by dissection under a magnifying glass. Glands were pooled and immediately rinsed three times in cold-PBS (137 mM NaCl, 1.45 mM NaH_2_PO_4_.H_2_O, 20.5 mM Na_2_HPO_4_, pH 7.2). They were then sonicated (Bandelin HD 2070, equipped with the MS 72 probe) three times for 15 seconds in 1.5 ml of PBS on ice, at 37% power, before concentration to a final volume of 100 µl with the Amicon Ultra filters (3 kDa threshold) and stored at −80°C until use in western blot. The remaining 17 ticks were then dissected under a magnifying glass and salivary glands, midgut, ovaries, synganglion and finally the rest of tick body (carcasse) were isolated, rinsed three times in cold-PBS, pooled according to organ and stored at −80°C until RNA extraction.

### Total RNA extraction

To evaluate gene expression, RNA extraction was performed on eggs, whole unfed larvae, nymphs, adult males and females, fed nymphs from the tick colony, and on tick organs from pre-fed field-collected females using the Nucleo Spin RNA (Macherey-Nagel) kit according to the manufacturer’s instructions. Tissue homogenization was first achieved with Precellys24 (Bertin instruments) in a lysis buffer containing β-mercaptoethanol with silicate beads for organs and metal beads (2.8mm of diameter) for whole ticks and eggs. RNA was eluted in 30 µl RNAse free water and RNA concentration was determined by measuring absorbance at 260 nm using a Nanodrop one (Thermo Fisher Scientific).

### Quantitative RT-PCR

To evaluate *IrSPI* expression at different tick life stages, RNA samples from one egg deposition, 120 unfed larvae, 20 unfed nymphs, 5 unfed males, and 5 unfed females from the tick colony were used. The temporal expression profile of *IrSPI* during feeding was evaluated in RNA samples from 20 unfed, 20 three-day, and 20 five-day fed nymphs from the colony after feeding on rabbits as previously described [68]. To evaluate *IrSPI* expression in different tick organs, pooled RNA samples from salivary glands, midgut, synganglion, ovaries, and carcasses of 17 females from the field that were pre-fed for five days on rabbits were used. Finally, the impact of tick infection with either transmitted or untransmitted bacteria was evaluated. For *B. birtlesii,* pooled salivary gland, gut, and ovary samples from 8 infected nymphs pre-fed for three days were used, while the same organs from 4 uninfected ticks were used as control. For *B. henselae*, 13 salivary gland, 14 gut, and 16 ovary pooled samples from infected females and non-infected controls pre-fed for four days were used. Finally, pooled RNA samples from 3 *E. coli*-infected females from the colony were analysed alongside 3 PBS-injected ticks or 3 uninjected ticks as controls.

Reverse transcription was performed with SuperScript III (Invitrogen) with 100 to 400 ng RNA per reaction. First strand cDNA synthesis was performed by using 2.5 µM Oligo dT primers in a total volume of 20 µl at 65°C for 5 min, 4°C for 3 min, 50°C for 50 min, and 70°C for 15 min. Following RNase H treatment (Invitrogen) at 37°C for 20 minutes, expression of *IrSPI, WecA (E. coli), defensin 6*, and *rps4* (a tick housekeeping gene) was evaluated by qPCR using the LightCycler 480 System (Roche). The SyBr green Master mix (Roche) added to 2 µl of cDNA, contained 6 µl of SyBr green vial 2, and 0.5 µM of both forward and reverse primers in a total volume of 12 µl. Previously described primers were used to detect *IrSPI* transcripts [5] while, for *defensin-6,* primers were designed according to the sequence published by Tonk *et al.* [9]: (F: 5’-TCGTCGTGATGATTGCGGGT-3’ and R: 5’-TCGTCGTGATGATTGCGGGT-3’). The following qPCR cycle conditions were used: 95°C 5 min; 95°C 10 s, 60°C 15 s, 72°C 15 s, for 45 cycles. Each sample was amplified in triplicate and results were analysed with Roche LightCycler 480 Software V1.5.0. Relative quantification of gene expression was calculated using the comparative Cp method [70]. *IrSPI* Cp were then normalised using the *I. ricinus rps4* gene [71] and expressed as fold change after ΔΔCp analysis. Analysis of *IrSPI* expression in wild female organs was performed using a plasmid containing *IrSPI* cDNA as a positive control, and *IrSPI* quantity was expressed as *IrSPI* cDNA copy number.

### *In situ* hybridization

*In situ* hybridization was performed as previously described [72]. Briefly, a 183-nucleotide long single-stranded DNA probe containing digoxygenin dNTP was prepared *via* asymmetric PCR (Roche Molecular Biochemicals) and the following primers: 5’-TGACTGAGACACAATGCAGA-3’ (forward) and 5’-TTCCGTACGGACATTCCCGC-3’ (reverse). The salivary glands from partially fed pathogen-free *I. ricinus* females engorged for eight days by means of a membrane feeding system [63] were dissected in ice-cold PBS and subsequently fixed in 4% paraformaldehyde (Sigma-Aldrich P6148) for 2 hours at room temperature. After three washes in PBST (PBS + 0.2% Triton), the tissues were treated with 25 µg/ml proteinase K for 10 minutes at RT. The reaction was stopped by adding PBST-glycine (0.2% of triton and 2 mg/ml glycine) for 5 minutes. Samples were re-fixed in 4% paraformaldehyde at RT and washed in PBST three times for 5 minutes. Hybridization was performed with either sense or antisense probe diluted ten times in hybridization solution (HS; 50% formamide, 5x SSC, 50 g/ml heparin, 0.1% Tween 20, and 100 g/ml salmon sperm DNA). The samples were then incubated in HS containing the representative probe for 30 hours at 48°C in a humidified chamber. After hybridization, tissues were washed sequentially in HS (12 hours, 48°C), HS-PBST (1:1; 5 min, RT) and PBST (5 min, RT). After blocking in 1% BSA in PBST for 20–30 min at RT, the tissues were immediately incubated with sheep anti-digoxigenin-AP (alkaline phosphatase; Roche Diagnostics, Germany) at a dilution of 1:1000 in PBST overnight at 4°C. Tissues were washed three times with PBST. Colorimetric reactions were developed by incubation with staining solution made from NBT/BCIP Ready-to-Use Tablets (Roche) according to the manufacturer’s protocol. Salivary glands were mounted in glycerol and examined under an optical microscope (Olympus BX53). Digital captured images were assembled and enhanced in Adobe Photoshop CS4.

### Anti-IrSPI serum production

Anti-IrSPI sera were produced against both the endogenous protein following tick bites in rabbits, and the recombinant protein in mice. Two hundred nymphs and around 2000 larvae from our *I. ricinus* colony were fed on New Zealand white rabbits (Charles River) until feeding completion (seven days) as previously described [68]. At day 17 post-infestation, rabbit was sacrificed and blood recovered by cardiac puncture. Five BALB/c mice were used to produce antibodies against IrSPI over a 90-day protocol. Three subcutaneous vaccinations (Days 0, 14, and 28) of 10 µg recombinant IrSPI were performed. Blood samples were collected retro-orbitally at 0, 14, 28, 42, and 56 days after the primary injection, and through final cardiac puncture at day 90. Blood from both mice and rabbits was stored at RT for 2 hours before two-step centrifugation at 6500g for 5 min to obtain clear serum that was stored at −20°C until used. Recombinant protein immunogenicity and anti-IrSPI murine antibody affinity were confirmed through ELISA experiments.

### Western blot analysis

Ten µl of both saliva and salivary gland extracts, as well as 1.5 µg of recombinant IrSPI protein as positive control, were analysed on 4-15% acrylamide gels under reducing conditions (Mini protean TGX stain free gels, Bio-Rad) followed by UV detection (ChemiDoc, Bio-Rad), or were immunoblotted with sera from mice or rabbits after IrSPI immunisation or tick infestation, respectively. Transfer to nitrocellulose membranes was performed using the transblot turbo (Bio-Rad) followed by saturation in TBS-5% milk for 1h at RT and incubation for 1h at RT with primary antibodies diluted in TBS-5% milk (1/1000 dilution for mice immune serum and 1/100 for rabbit serum). After 3 washes in TBS-5% milk, antibody binding was detected with the BCIP/NBT Color development system (Roche), using either anti-mouse or anti-rabbit alkaline phosphatase-conjugated antibodies (A3562 or A3687, respectively; Sigma-Aldrich) diluted 1:7500 in TBS-5%. Results were analysed with the ChemiDoc instrument (Bio-Rad) and the accompanying ImageLab 1.5 software (Bio-Rad).

### Thrombin generation, clot waveform, and fibrinolysis assays

Thrombin generation assays (TGA) were adapted from the calibrated automated thrombogram (CAT) as described by Jourdi *et al*. [73]. Briefly, 80 μl calibrated platelet-poor plasma (PPP, Cryopep, Montpellier, France) with or without 0.8 µM refolded IrSPI, and 20 μl PPP-Reagent (Stago, Asnières– sur-Seine, France) were dispensed into wells of flat-bottomed microtiter plates. After 5 min of preheating at 37°C, thrombin generation was initiated by automated injection of 20 μL of a substrate solution and calcium mixture (FluCa, Stago). Progress of Z-Gly-Gly-Arg-7-amino-4-methylcoumarin hydrolysis was recorded every 10 seconds by a thermostated Tecan Infinite M200 Pro (excitation 340 nm, emission 440 nm). Fluorescence measurements were analysed using GraphPad Prism software. Lag time, time to peak (TTP), maximum amount of thrombin formed (peak height), and endogenous thrombin potential (ETP) were computed using the area under curve method. Clot waveform assays were simultaneously performed by utilising the Tecan reader’s capacity to simultaneously record fluorescence and A_405_ as described by Jourdi *et al.* [74]. Data were analysed using GraphPad Prism software to estimate the lag time of clot formation, defined as the time needed to reach 15% of maximum turbidity. Fibrinolysis was measured by the clot waveform assay except that 6 nM of tPA (Actilyse, Boehringer Ingelheim, Germany) was added to the PPP prior to triggering clot formation. Time to half-lysis was defined as the time needed to halve the maximum turbidity. Obtained results were then compared between positive controls, negative controls (refolding buffer), and samples treated with IrSPI.

### IrSPI impact on endothelial cell apoptosis

The mature human skin microvascular endothelial cell line, HSkMEC (Human Skin Microvascular Endothelial Cell line) [75], was cultivated in 24-well plates in complete OptiMEM® medium (OptiMEM® with 2% FBS, 0.5 µg/mL fungizone, and 40 µg/mL gentamicin, Gibco). After 24 hours, medium was replaced by 1 µM of recombinant IrSPI that had been refolded or not, or by their cognate controls, that is, refolding buffer or complete OptiMEM® medium, respectively. Cells were then incubated for 40 h at 37°C. At 24 h, staurosporin (1µM) was added to the corresponding wells as a positive apoptosis control. Cells were detached by trypsinisation, washed, labelled with Annexin-V-FITC and propidium iodide according to the manufacturer’s instructions (FITC Annexin V kit and Dead Cell Apoptosis Kit, Invitrogen), and analysed by cytofluorimetry with the LSR Fortessa X20 instrument (Becton Dickinson).

### IrSPI impact on host angiogenesis

The potential cytotoxicity of IrSPI was initially evaluated on HSkMEC cells. After 24 h of culture in 96-well plates in complete OptiMEM® medium, the medium was replaced by refolded or untreated IrSPI at 2, 1, 0.5, 0.25, 0.125, or 0.0625 µM, or by the corresponding controls; namely, refolding buffer or complete OptiMEM® medium. After 48 h of culture, each supernatant was removed and replaced by a solution of Alamar Blue^TM^ (100 µl, at 1/22 dilution in OptiMEM® medium, Invitrogen). Cell toxicity was assessed after 3 h of incubation by measuring fluorescence intensity using the Victor 3V spectrofluorometric multiwell plate reader (Perkin Elmer; excitation at 560 nm; emission at 605 nm). The ability of IrSPI, refolded or not, to modulate host angiogenesis was then evaluated on two human endothelial cell lines, HSkMEC and HEPC-CB1 (Human Endothelial Progenitor Cells-Cord Blood1), a human progenitor endothelial cell line [76]. To coat plates, 50 µl of Matrigel^TM^ (50% dilution in OptiMEM® medium) was dispensed into 96-well microplates and incubated for 30 min at 37°C to allow Matrigel^TM^ polymerization. Cells (1.5 × 10^5^ cells/ml) were resuspended in 100 µl OptiMEM® containing either 1 or 0.5 µM of the recombinant protein, refolded or not, or, as controls, the refolding buffer or culture medium, and were then seeded into the Matrigel^TM^ matrix-containing wells. Plates were incubated in a videomicroscope chamber at 37°C with 5% CO_2_. Pseudo vessel formation was visualised for 9.5 h with an inverted microscope equipped with a CCD camera (Axio Observer Z1, Zeiss, Le Pecq, France), with acquisitions every 30 min *via* the Zen software (Zeiss). Angiogenesis was quantified with the ImageJ software (with Angiogenesis Analyzer) according to several parameters: number of segments and length, number of meshes, and number of junctions [77].

### Splenocyte proliferation assay

Spleens of three OF1 mice (8 weeks of age) were gently crushed in individual cell strainers (100 µm). After 2 washes in complete medium (RPMI 1640 (Gibco) with 5% sodium pyruvate, 1% penicillin/streptomycin (Invitrogen), 1% 200 mM L-glutamine (Gibco), and 10% heat-inactivated FCS (Invitrogen)), cells were centrifuged for 5 min at 1400 rpm and incubated at 37°C for 2 min in sterile RBC lysis solution (Invitrogen). Lysis was arrested by adding 30 ml PBS, and cells were pelleted and resuspended in 2 ml complete medium. Cells were then labelled with CFSE (10 µM, Molecular Probes) for 5 minutes at RT in the dark, and quenched by adding 5 volumes of cold complete medium. CFSE-labelled cells were maintained on ice for 5 min and then washed twice in complete medium. The proliferation of mitogen-stimulated cells was then measured in the presence or absence of IrSPI, in triplicate wells for each condition. For each mouse, CFSE-labelled cells (5 × 10^5^ per well in 50 µl) were dispensed in 96-well U-bottomed plates. Recombinant IrSPI (2 µg per well in 50 µl complete medium) or medium alone was added and cells were incubated at 37°C in 5% CO_2_ for 2 hours. Concanavalin A (ConA) (2 µg/ml in 100 µl of complete medium containing 0.1 mM ß-mercaptoethanol) was then added. Single wells containing CFSE-labelled and unlabelled cells were included for isotype controls and for spectral compensation for CFSE and the viability label. After 3 days of culture (37°C in 5% CO_2_), supernatants (150 µl/ well) were collected and stored at −80°C for subsequent analysis of cytokine expression and cells were harvested for phenotypic labelling and flow cytometry.

For phenotypic labelling, all antibodies were purchased from eBiosciences and diluted in PBS containing 0.5% bovine serum albumin and 2 mM EDTA (PBS–BSA–EDTA). To block Fc receptors, cells were resuspended in 50 µl of anti-mouse-CD16/32 (0.5 µg/well) and incubated for 15 min at 4°C. Cells were then resuspended in a cocktail of PE-anti-CD4 and APC-anti-CD8 (both at 0.125 µg/well) and eFluor 450-anti-CD19 and APC-eFluor 780-anti-CD3e (both at 0.5 µg/well). For isotypic controls, the same concentration of the cognate isotypic antibodies was used. After 30 min, cells were washed and labelled for 30 min with a viability marker (LIVE/DEAD® Fixable Aqua Dead Cell Stain Kit, Molecular Probes) according to the manufacturer’s recommendations. Cells were washed and fixed with 150 µl of 1% paraformaldehyde in PBS. Flow cytometry was performed using the FACSCanto II cytometer (BD Biosciences). Data were acquired and analysed with FACSDiva (BD Biosciences) and FlowJo software (Tree Star), respectively. Live cells, distinguished by referring to dot plots of Aqua staining versus side scatter, were gated as B or T lymphocytes on the basis of CD19 or CD3e staining, respectively, and T cells were further distinguished for CD4 or CD8 expression. The CFSE staining of each subset was then examined.

### Macrophage stimulation assay

Macrophages were isolated from bone marrow of three 20-week-old BALB/cJRj mice. Mice were sacrificed by cervical dislocation and femurs and tibias were harvested and kept on ice in PBS-A (without Mg^2+^ and Ca^2+^). Bone marrow was flushed out from cleaned bones without muscle with cold PBS-A using a 10 ml syringe and 25-gauge needle into a sterile 50 ml conical tube placed on ice. Bone marrow cells were centrifuged for 10 min at 4°C and 300g, the pellet was resuspended in 500 µl of ice-cold PBS-A. Five ml of Gey buffer was added drop by drop to the cell suspension to lyse contaminating red blood cells. After 10 minutes of incubation at 4°C, volume was completed to 50 ml with cold PBS-A and the cell suspension centrifuged for 10 minutes at 4°C and 300g. The cell pellet was carefully resuspended in 1 ml of complete medium to dissociate aggregates. The cell suspension was diluted in complete medium before cell counting using Malassez microscopy counting slides. Suspensions of 1.5 × 10^6^ cells/ml were cultivated at 37°C with 7.5% of CO_2_ in Petri dishes (Petri Greiner ref: 664161) using complete DMEM medium (Pan DMEM (Biotech P04-03588) with 15% FCS and 4.5 g/l glucose, 50 µg/ml penicillin and streptomycin, 10 mM HEPES, 50 µM β−ME) supplemented with 50 ng/ml M-CSF-1 (rm M-CSF-1, ref 12343115, ImmunoTools). Fresh medium was added after three days of culture. At day six, supernatants were discarded into Falcon tubes and adherent macrophages were harvested by gentle flushing following a 30 min incubation at 37°C in PBS-A containing 25 mM EDTA. Individual cell suspensions (1 for each of 3 mice) were then transferred to 12-well plates by using 10 ml of PBS-A at 2 × 10^6^ cells/wells. Four culture conditions were tested for 24 h in complete medium: with or without IrSPI (10 µg/ml), and with or without stimulation cocktail (0.5 µg/ml LPS and 100 U/ml IFN-γ). Respective controls were complete medium or complete medium with stimulation cocktail.

### Cytokine profile analysis

The impact of IrSPI on cytokine production was evaluated in splenocytes (with or without ConA stimulation) and in LPS and IFN-γ-activated macrophages with the Luminex Procarta PPX-01-S26088EX kit (Invitrogen) targeting mouse Th1 (IL2, IL-1β, IFN-γ, TNFα, IL12p70, IL-18), Th2 (IL4, IL5, IL6, IL-10, IL-13, IL9), Th17 (IL-17, IL-23, IL-27), and Th22 (IL-22) cytokines, chemokines (Eotaxin, Groα-KC, IP10, MIP-1α, MIP-1β, MIP2, MCP1, MCP3, RANTES), and one colony-stimulating factor (GM-CSF), according to the manufacturers’ instructions. Supernatant (50 µl) from splenocytes cultured with or without mitogen stimulation, and with or without IrSPI, and 100 µl of supernatant from activated macrophages cultivated with or without IrSPI, as described above, were used. Each supernatant was incubated for 1.5 h with magnetic beads coated with antibodies directed against the targeted cytokines. The beads with immobilised cytokines were then successively incubated with biotinylated anti-cytokine antibodies and streptavidin-PE, and analysed using the Luminex^TM^ 200 system (Bio-Rad).

### Statistical analysis

GraphPad Prism version 7.8 was used to perform ANCOVA test for inhibition assay linear regression analysis, Student’s t-test to evaluate the significance of IrSPI expression and angiogenesis parameters, as well as splenocyte proliferation differences in response to IrSPI exposure. P-values < 0.05 were considered as significant (labelled with *) whereas more significant results with P-values < 0.01 were indicated with **.

Cytokine/chemokine Luminex expression analysis was performed for each of 3 mice in triplicate, and in duplicate for splenocytes and macrophages. ANOVA analysis was used to evaluate the variability between triplicates or duplicates and between mice for each measured molecule, with and without stimulation. The variable “presence or absence of IrSPI” and the variable mouse were used as fixed and random effect, respectively. The influence of the protein on the variable “cell response” was expressed as P values (P<0.01).

### Software

NetNGlyc 1.0 and NetPhos 3.1 servers (DTU Bioinformatics) and Phyre 2 software [78] were used in order to predict post-translational modifications and secondary structures of IrSPI, respectively.

## Acknowledgments

The authors gratefully acknowledge Robert Menard for his precious help for editing this paper. We thank Martine Cote, Evelyne Le-Naour and Lisa Fourniol for tick collection, tick rearing and technical assistance. We also thank the CBM P@CYFIC platform (Orleans, France) for video-microscopy of angiogenesis experiments and Dr Claudine Kieda for the establishment of the human endothelial cell lines. We also thank Sara Moutailler for technical advice in qPCR experiments and Bruno Baron from the *Plateforme de Biophysique moléculaire* (Institut Pasteur) for help with CD experiments and data analysis. We also thank the “*Tiques et Maladies à Tiques*” group (*REID-Réseau Ecologie des Interactions Durables*) for stimulating discussion.

## Author contributions

AAB and SIB conceived the experiments. AAB, SIB, JR, LS, CG, FF, SB, BLB, SSBB, AR and EP designed and performed experiments. AAB, SIB, JR, LS, SB, BLB, CG, and MM analysed the data. AAB, SIB, JR, LS, CG, FF, SB, BLB, EP, and MM wrote the paper.

## Supporting information

**S1 Fig. IrSPI MALDI TOF/TOF analysis.** A volume of 1 µl at 0.075 mg/mL was deposited on a MTP 384 ground steel target plate with 1 µL2,5-DHB at 25 mg/ml prepared in 50% acetonitrile, 0.1% trifluoroacetic acid in water as a matrix solution. Data were acquired using Flexcontrol software (Bruker-Daltonics, Germany) and shots were recorded in positive ion linear mode.

**S2 Fig. Circular dichroism analyses of recombinant IrSPI at 0.075mg/ml**. Experiments were performed in the far-UV using an Aviv CD spectrometer model 215 equipped with a water-cooled Peltier unit. Spectra were recorded in a cell width of 1-mm path length in 200µl in far-UV (between 190 to 270 nm) at 25°C. IrSPI and its cognate buffer were successively screened ten times to produce an averaged spectrum, which was corrected using buffer baselines. Data were normalised to residue molar absorption measured in mdeg (M^−1^ cm^−1^) and expressed as delta epsilon (Δε). BestSel website algorithm treatment was used to predict secondary structures (Micsonai et al. BeStSel: a web server for accurate protein secondary structure prediction and fold recognition from the circular dichroism spectra Nucleic Acids Research, Volume 46, Issue W1, 2 July 2018, Pages W315–W322).

**S3 Fig. Analytical ultracentrifugation analysis of the recombinant IrSPI at 0.075µg/µl in epoxy double sector cell**. The measurement cell was equilibrated for 1.5 h at 20°C in a four-hole AN60-Ti rotor. The sample was spun for 10 h and 400 scans for both Rayleigh interference and absorbance at 280 nm were recorded. Absorbance data were collected at a constant radial step size of 0.001 cm. Partial specific volume of 0.70 and the extinction coefficient ε at 280 nm of 14350 L.mol^-1^.cm^-1^ were theoretically calculated at 20°C from the IrSPI amino acid sequence using Sednterp software (Spin Analytical, USA). As a control, the PBS (-Ca^2+^ and -Mg^2+^) density ρ and viscosity η of 1.0053 g.mL^-1^ and 1.018 cP respectively were also determined with Sednterp at 20°C. Data were analysed with Sedfit.

**S1 Table:**
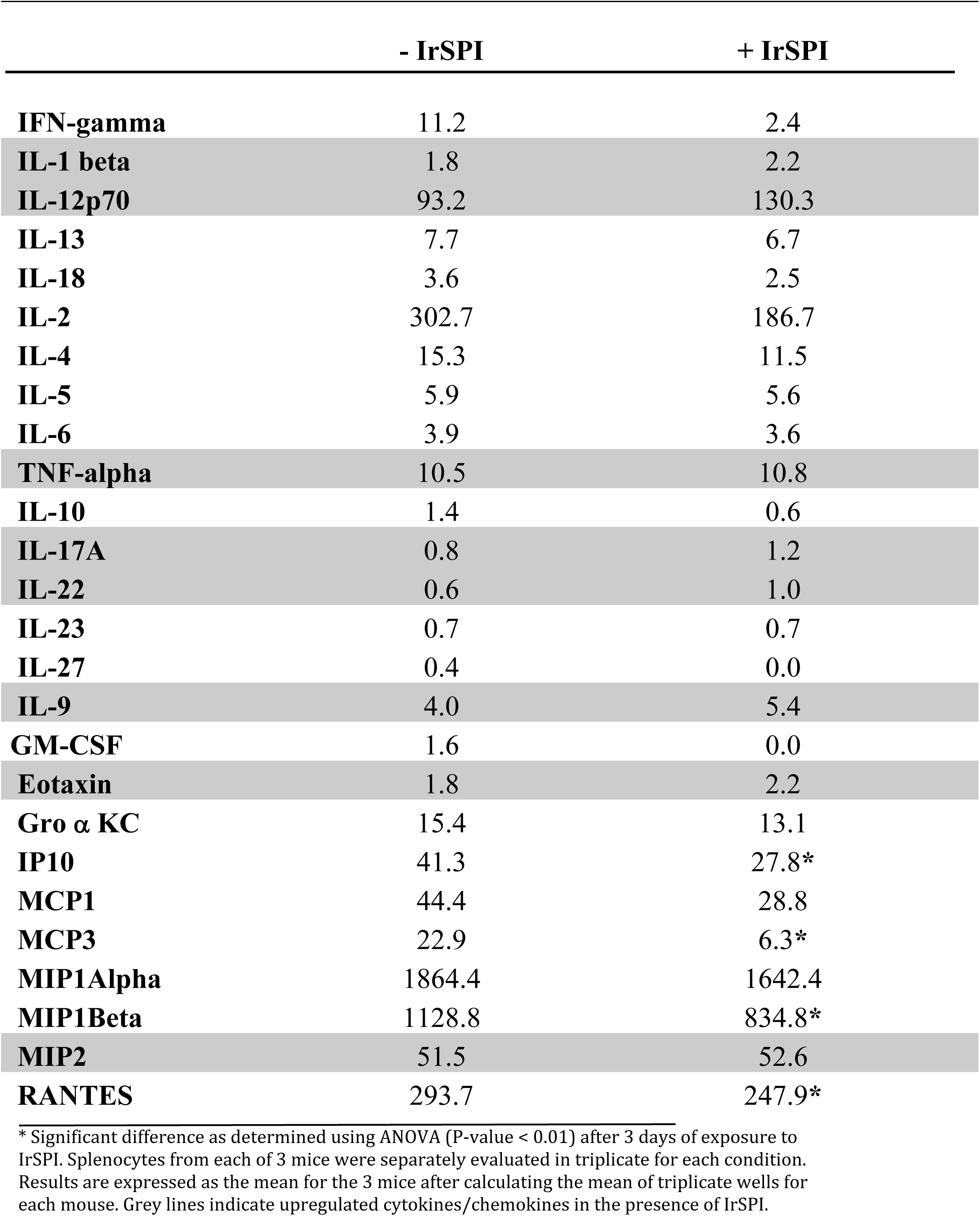
Cytokine and chemokine concentrations (pg/ml) in splenocyte supernatants in the presence or absence of IrSPI without ConA stimulation and normalised to standards

**S2 Table:** Cytokine and chemokine concentrations (pg/ml) in splenocyte supernatants in the presence or absence of IrSPI after ConA stimulation and normalised to standards.

|  | - IrSPI | + IrSPI |
| --- | --- | --- |
| **IFN-gamma** | 7744.2 | 3849.7* |
| **IL-1 beta** | 14.3 | 7.9* |
| **IL-12p70** | 147.2 | 141.8 |
| **IL-13** | 1000.1 | 248.9* |
| **IL-18** | 157.1 | 84.8* |
| **IL-2** | 867.3 | 1293.8* |
| **IL-4** | 719.9 | 547.2 |
| **IL-5** | 48.3 | 20.1 |
| **IL-6** | 134.8 | 60.9* |
| **TNF-alpha** | 372.1 | 200.5* |
| **IL-10** | 24.0 | 19.7 |
| **IL-17A** | 404.4 | 224.1 |
| **IL-22** | 15.4 | 5.2 |
| **IL-23** | 1.3 | 1.4 |
| **IL-27** | 1.4 | 0.1 |
| **IL-9** | 116.7 | 59.7* |
| **GM-CSF** | 537.2 | 150.7* |
| **Eotaxin** | 6.8 | 5.3* |
| **Gro α KC** | 53.3 | 45.9 |
| **IP10** | 1448.3 | 774.4* |
| **MCP1** | 306.4 | 276.7 |
| **MCP3** | 248.3 | 202.7 |
| **MIP1Alpha** | 9645.8 | 8942.3 |
| **MIP1Beta** | 7711.5 | 6328.5* |
| **MIP2** | 135.0 | 109.9 |
| **RANTES** | 1633.1 | 1013.7* |

\* Significant difference as determined using ANOVA (P-value < 0.01) after 3 days of exposure to IrSPI. Splenocytes from each of 3 mice were separately evaluated in triplicate for each condition. Results are expressed as the mean for the 3 mice after calculating the mean of triplicate wells for each mouse. Grey lines indicate upregulated cytokines/chemokines in the presence of IrSPI.

**S3 Table:**
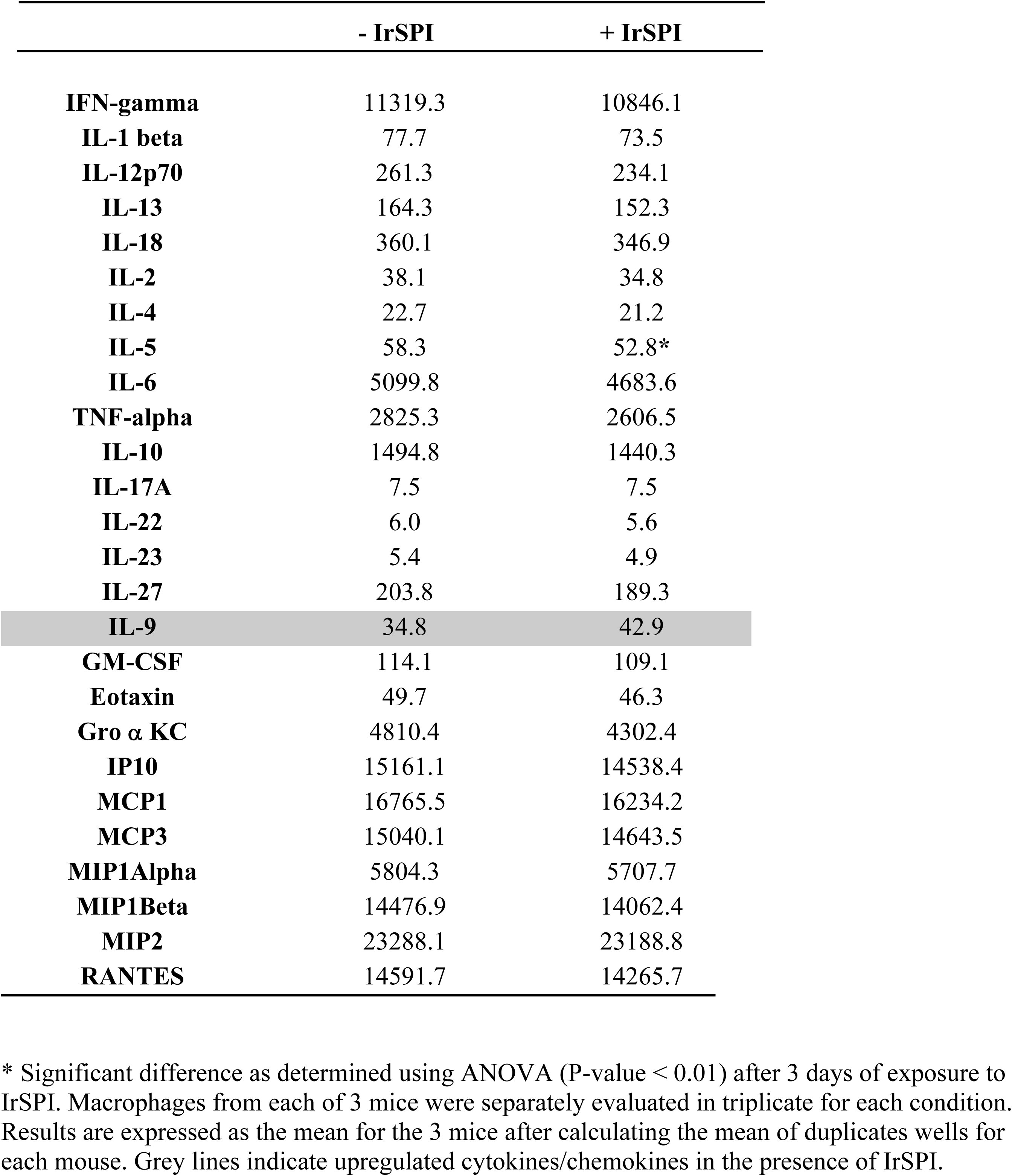
Cytokine and chemokine concentrations (pg/ml) in supernatants of activated macrophages in the presence or absence of IrSPI and normalised to standards

|  | - IrSPI | + IrSPI |
| --- | --- | --- |
| **IFN-gamma** | 11319.3 | 10846.1 |
| **IL-1 beta** | 77.7 | 73.5 |
| **IL-12p70** | 261.3 | 234.1 |
| **IL-13** | 164.3 | 152.3 |
| **IL-18** | 360.1 | 346.9 |
| **IL-2** | 38.1 | 34.8 |
| **IL-4** | 22.7 | 21.2 |
| **IL-5** | 58.3 | 52.8* |
| **IL-6** | 5099.8 | 4683.6 |
| **TNF-alpha** | 2825.3 | 2606.5 |
| **IL-10** | 1494.8 | 1440.3 |
| **IL-17A** | 7.5 | 7.5 |
| **IL-22** | 6.0 | 5.6 |
| **IL-23** | 5.4 | 4.9 |
| **IL-27** | 203.8 | 189.3 |
| **IL-9** | 34.8 | 42.9 |
| **GM-CSF** | 114.1 | 109.1 |
| **Eotaxin** | 49.7 | 46.3 |
| **Gro α KC** | 4810.4 | 4302.4 |
| **IP10** | 15161.1 | 14538.4 |
| **MCP1** | 16765.5 | 16234.2 |
| **MCP3** | 15040.1 | 14643.5 |
| **MIP1Alpha** | 5804.3 | 5707.7 |
| **MIP1Beta** | 14476.9 | 14062.4 |
| **MIP2** | 23288.1 | 23188.8 |
| **RANTES** | 14591.7 | 14265.7 |

\* Significant difference as determined using ANOVA (P-value < 0.01) after 3 days of exposure to IrSPI. Macrophages from each of 3 mice were separately evaluated in triplicate for each condition. Results are expressed as the mean for the 3 mice after calculating the mean of duplicates wells for each mouse. Grey lines indicate upregulated cytokines/chemokines in the presence of IrSPI.

## References

1. de la Fuente J, Estrada-Pena A, Venzal JM, Kocan KM, Sonenshine DE. Overview: Ticks as vectors of pathogens that cause disease in humans and animals. Front Biosci. 2008;13:6938–46. Epub 2008/05/30. PubMed PMID: 18508706.

2. Rizzoli A, Silaghi C, Obiegala A, Rudolf I, Hubalek Z, Foldvari G, et al. Ixodes ricinus and Its Transmitted Pathogens in Urban and Peri-Urban Areas in Europe: New Hazards and Relevance for Public Health. Front Public Health. 2014;2:251. doi: 10.3389/fpubh.2014.00251. PubMed PMID: 25520947; PubMed Central PMCID: PMCPMC4248671.

3. Simo L, Kazimirova M, Richardson J, Bonnet SI. The Essential Role of Tick Salivary Glands and Saliva in Tick Feeding and Pathogen Transmission. Front Cell Infect Microbiol. 2017;7:281. Epub 2017/07/12. doi: 10.3389/fcimb.2017.00281. PubMed PMID: 28690983; PubMed Central PMCID: PMCPMC5479950.

4. de la Fuente J, Antunes S, Bonnet S, Cabezas-Cruz A, Domingos AG, Estrada-Pena A, et al. Tick-Pathogen Interactions and Vector Competence: Identification of Molecular Drivers for Tick-Borne Diseases. Front Cell Infect Microbiol. 2017;7:114. doi: 10.3389/fcimb.2017.00114. PubMed PMID: 28439499; PubMed Central PMCID: PMCPMC5383669.

5. Liu XY, de la Fuente J, Cote M, Galindo RC, Moutailler S, Vayssier-Taussat M, et al. IrSPI, a tick serine protease inhibitor involved in tick feeding and Bartonella henselae infection. PLoS Negl Trop Dis. 2014;8(7):e2993. Epub 2014/07/25. doi: 10.1371/journal.pntd.0002993. PubMed PMID: 25057911; PubMed Central PMCID: PMCPMC4109860.

6. Cotte V, Bonnet S, Le Rhun D, Le Naour E, Chauvin A, Boulouis HJ, et al. Transmission of Bartonella henselae by Ixodes ricinus. Emerg Infect Dis. 2008;14(7):1074–80. doi: 10.3201/eid1407.071110. PubMed PMID: 18598628; PubMed Central PMCID: PMCPMC2600320.

7. Blisnick AA, Foulon T, Bonnet SI. Serine Protease Inhibitors in Ticks: An Overview of Their Role in Tick Biology and Tick-Borne Pathogen Transmission. Front Cell Infect Microbiol. 2017;7:199. Epub 2017/06/08. doi: 10.3389/fcimb.2017.00199. PubMed PMID: 28589099; PubMed Central PMCID: PMCPMC5438962.

8. Paul RE, Cote M, Le Naour E, Bonnet SI. Environmental factors influencing tick densities over seven years in a French suburban forest. Parasit Vectors. 2016;9(1):309. Epub 2016/05/29. doi: 10.1186/s13071-016-1591-5. PubMed PMID: 27234215; PubMed Central PMCID: PMCPMC4884405.

9. Tonk M, Cabezas-Cruz A, Valdes JJ, Rego RO, Grubhoffer L, Estrada-Pena A, et al. Ixodes ricinus defensins attack distantly-related pathogens. Dev Comp Immunol. 2015;53(2):358–65. Epub 2015/08/10. doi: 10.1016/j.dci.2015.08.001. PubMed PMID: 26255244.

10. Simo L, Zitnan D, Park Y. Neural control of salivary glands in ixodid ticks. J Insect Physiol. 2012;58(4):459–66. Epub 2011/11/29. doi: 10.1016/j.jinsphys.2011.11.006. PubMed PMID: 22119563; PubMed Central PMCID: PMCPMC3295888.

11. Kotal J, Langhansova H, Lieskovska J, Andersen JF, Francischetti IM, Chavakis T, et al. Modulation of host immunity by tick saliva. J Proteomics. 2015;128:58–68. Epub 2015/07/21. doi: 10.1016/j.jprot.2015.07.005. PubMed PMID: 26189360; PubMed Central PMCID: PMCPMC4619117.

12. Francischetti IM, Mather TN, Ribeiro JM. Tick saliva is a potent inhibitor of endothelial cell proliferation and angiogenesis. Thromb Haemost. 2005;94(1):167–74. Epub 2005/08/23. doi: 10.1160/TH04-09-0566. PubMed PMID: 16113800; PubMed Central PMCID: PMCPMC2893037.

13. Mans BJ, Louw AI, Neitz AW. Evolution of hematophagy in ticks: common origins for blood coagulation and platelet aggregation inhibitors from soft ticks of the genus Ornithodoros. Mol Biol Evol. 2002;19(10):1695–705. doi: 10.1093/oxfordjournals.molbev.a003992. PubMed PMID: 12270896.

14. Alarcon-Chaidez FJ, Muller-Doblies UU, Wikel S. Characterization of a recombinant immunomodulatory protein from the salivary glands of Dermacentor andersoni. Parasite Immunol. 2003;25(2):69–77. PubMed PMID: 12791102.

15. Konnai S, Nishikado H, Yamada S, Imamura S, Ito T, Onuma M, et al. Molecular identification and expression analysis of lipocalins from blood feeding taiga tick, Ixodes persulcatus Schulze. Exp Parasitol. 2011;127(2):467–74. doi: 10.1016/j.exppara.2010.10.002. PubMed PMID: 21036169.

16. Garcia-Fernandez R, Perbandt M, Rehders D, Ziegelmuller P, Piganeau N, Hahn U, et al. Three-dimensional Structure of a Kunitz-type Inhibitor in Complex with an Elastase-like Enzyme. J Biol Chem. 2015;290(22):14154–65. doi: 10.1074/jbc.M115.647586. PubMed PMID: 25878249; PubMed Central PMCID: PMCPMC4447985.

17. Wilgus TA, Roy S, McDaniel JC. Neutrophils and Wound Repair: Positive Actions and Negative Reactions. Adv Wound Care (New Rochelle). 2013;2(7):379–88. doi: 10.1089/wound.2012.0383. PubMed PMID: 24527354; PubMed Central PMCID: PMCPMC3763227.

18. Shah D, Mital K. The Role of Trypsin:Chymotrypsin in Tissue Repair. Adv Ther. 2018;35(1):31–42. doi: 10.1007/s12325-017-0648-y. PubMed PMID: 29209994; PubMed Central PMCID: PMCPMC5778189.

19. Valdes JJ, Schwarz A, Cabeza de Vaca I, Calvo E, Pedra JH, Guallar V, et al. Tryptogalinin is a tick Kunitz serine protease inhibitor with a unique intrinsic disorder. PLoS One. 2013;8(5):e62562. Epub 2013/05/10. doi: 10.1371/journal.pone.0062562. PubMed PMID: 23658744; PubMed Central PMCID: PMCPMC3643938.

20. Liu XY, Bonnet SI. Hard tick factors implicated in pathogen transmission. PLoS Negl Trop Dis. 2014;8(1):e2566. doi: 10.1371/journal.pntd.0002566. PubMed PMID: 24498444; PubMed Central PMCID: PMCPMC3907338.

21. Kotsyfakis M, Schwarz A, Erhart J, Ribeiro JM. Tissue- and time-dependent transcription in Ixodes ricinus salivary glands and midguts when blood feeding on the vertebrate host. Sci Rep. 2015;5:9103. Epub 2015/03/15. doi: 10.1038/srep09103. PubMed PMID: 25765539; PubMed Central PMCID: PMCPMC4357865.

22. Chmelar J, Anderson JM, Mu J, Jochim RC, Valenzuela JG, Kopecky J. Insight into the sialome of the castor bean tick, Ixodes ricinus. BMC Genomics. 2008;9:233. Epub 2008/05/21. doi: 10.1186/1471-2164-9-233. PubMed PMID: 18489795; PubMed Central PMCID: PMCPMC2410133.

23. Dai SX, Zhang AD, Huang JF. Evolution, expansion and expression of the Kunitz/BPTI gene family associated with long-term blood feeding in Ixodes Scapularis. BMC Evol Biol. 2012;12:4. Epub 2012/01/17. doi: 10.1186/1471-2148-12-4. PubMed PMID: 22244187; PubMed Central PMCID: PMCPMC3273431.

24. Islam MK, Tsuji N, Miyoshi T, Alim MA, Huang X, Hatta T, et al. The Kunitz-like modulatory protein haemangin is vital for hard tick blood-feeding success. PLoS Pathog. 2009;5(7):e1000497. Epub 2009/07/14. doi: 10.1371/journal.ppat.1000497. PubMed PMID: 19593376; PubMed Central PMCID: PMCPMC2701603.

25. Soares TS, Oliveira F, Torquato RJ, Sasaki SD, Araujo MS, Paschoalin T, et al. BmTI-A, a Kunitz type inhibitor from Rhipicephalus microplus able to interfere in vessel formation. Vet Parasitol. 2016;219:44–52. Epub 2016/02/28. doi: 10.1016/j.vetpar.2016.01.021. PubMed PMID: 26921038.

26. Lima CA, Torquato RJ, Sasaki SD, Justo GZ, Tanaka AS. Biochemical characterization of a Kunitz type inhibitor similar to dendrotoxins produced by Rhipicephalus (Boophilus) microplus (Acari: Ixodidae) hemocytes. Vet Parasitol. 2010;167(2-4):279–87. Epub 2009/10/16. doi: 10.1016/j.vetpar.2009.09.030. PubMed PMID: 19828254.

27. Drewes CC, Dias RY, Hebeda CB, Simons SM, Barreto SA, Ferreira JM, Jr., et al. Actions of the Kunitz-type serine protease inhibitor Amblyomin-X on VEGF-A-induced angiogenesis. Toxicon. 2012;60(3):333–40. doi: 10.1016/j.toxicon.2012.04.349. PubMed PMID: 22575283.

28. Davie EW, Fujikawa K, Kurachi K, Kisiel W. The role of serine proteases in the blood coagulation cascade. Adv Enzymol Relat Areas Mol Biol. 1979;48:277–318. PubMed PMID: 367103.

29. Mejri N, Franscini N, Rutti B, Brossard M. Th2 polarization of the immune response of BALB/c mice to Ixodes ricinus instars, importance of several antigens in activation of specific Th2 subpopulations. Parasite Immunol. 2001;23(2):61–9. PubMed PMID: 11240897.

30. Ramachandra RN, Wikel SK. Modulation of host-immune responses by ticks (Acari: Ixodidae): effect of salivary gland extracts on host macrophages and lymphocyte cytokine production. J Med Entomol. 1992;29(5):818–26. Epub 1992/09/01. PubMed PMID: 1404261.

31. Rodrigues V, Fernandez B, Vercoutere A, Chamayou L, Andersen A, Vigy O, et al. Immunomodulatory Effects of Amblyomma variegatum Saliva on Bovine Cells: Characterization of Cellular Responses and Identification of Molecular Determinants. Front Cell Infect Microbiol. 2017;7:521. Epub 2018/01/23. doi: 10.3389/fcimb.2017.00521. PubMed PMID: 29354598; PubMed Central PMCID: PMCPMC5759025.

32. Urioste S, Hall LR, Telford SR, 3rd, Titus RG. Saliva of the Lyme disease vector, Ixodes dammini, blocks cell activation by a nonprostaglandin E2-dependent mechanism. J Exp Med. 1994;180(3):1077–85. PubMed PMID: 8064226; PubMed Central PMCID: PMCPMC2191645.

33. Chang HC, Sehra S, Goswami R, Yao W, Yu Q, Stritesky GL, et al. The transcription factor PU.1 is required for the development of IL-9-producing T cells and allergic inflammation. Nat Immunol. 2010;11(6):527–34. doi: 10.1038/ni.1867. PubMed PMID: 20431622; PubMed Central PMCID: PMCPMC3136246.

34. Anguita J, Ramamoorthi N, Hovius JW, Das S, Thomas V, Persinski R, et al. Salp15, an ixodes scapularis salivary protein, inhibits CD4(+) T cell activation. Immunity. 2002;16(6):849–59. Epub 2002/07/18. PubMed PMID: 12121666.

35. Leboulle G, Crippa M, Decrem Y, Mejri N, Brossard M, Bollen A, et al. Characterization of a novel salivary immunosuppressive protein from Ixodes ricinus ticks. J Biol Chem. 2002;277(12):10083–9. Epub 2002/01/17. doi: 10.1074/jbc.M111391200. PubMed PMID: 11792703.

36. Skallova A, Iezzi G, Ampenberger F, Kopf M, Kopecky J. Tick saliva inhibits dendritic cell migration, maturation, and function while promoting development of Th2 responses. J Immunol. 2008;180(9):6186–92. PubMed PMID: 18424740.

37. Taub DD, Turcovski-Corrales SM, Key ML, Longo DL, Murphy WJ. Chemokines and T lymphocyte activation: I. Beta chemokines costimulate human T lymphocyte activation in vitro. J Immunol. 1996;156(6):2095–103. Epub 1996/03/15. PubMed PMID: 8690897.

38. Elyaman W, Bradshaw EM, Uyttenhove C, Dardalhon V, Awasthi A, Imitola J, et al. IL-9 induces differentiation of TH17 cells and enhances function of FoxP3+ natural regulatory T cells. Proc Natl Acad Sci U S A. 2009;106(31):12885–90. doi: 10.1073/pnas.0812530106. PubMed PMID: 19433802; PubMed Central PMCID: PMCPMC2722314.

39. de Waal Malefyt R, Abrams JS, Zurawski SM, Lecron JC, Mohan-Peterson S, Sanjanwala B, et al. Differential regulation of IL-13 and IL-4 production by human CD8+ and CD4+ Th0, Th1 and Th2 T cell clones and EBV-transformed B cells. Int Immunol. 1995;7(9):1405–16. PubMed PMID: 7495748.

40. Kovar L, Kopecky J, Rihova B. Salivary gland extract from Ixodes ricinus tick modulates the host immune response towards the Th2 cytokine profile. Parasitol Res. 2002;88(12):1066–72. doi: 10.1007/s00436-002-0714-4. PubMed PMID: 12444457.

41. Mejri N, Brossard M. Splenic dendritic cells pulsed with Ixodes ricinus tick saliva prime naive CD4+T to induce Th2 cell differentiation in vitro and in vivo. Int Immunol. 2007;19(4):535–43. doi: 10.1093/intimm/dxm019. PubMed PMID: 17344202.

42. Ganapamo F, Rutti B, Brossard M. In vitro production of interleukin-4 and interferon-gamma by lymph node cells from BALB/c mice infested with nymphal Ixodes ricinus ticks. Immunology. 1995;85(1):120–4. PubMed PMID: 7635513; PubMed Central PMCID: PMCPMC1384034.

43. Zeidner N, Dreitz M, Belasco D, Fish D. Suppression of acute Ixodes scapularis-induced Borrelia burgdorferi infection using tumor necrosis factor-alpha, interleukin-2, and interferon-gamma. J Infect Dis. 1996;173(1):187–95. PubMed PMID: 8537658.

44. Thelen M, Stein JV. How chemokines invite leukocytes to dance. Nat Immunol. 2008;9(9):953–9. doi: 10.1038/ni.f.207. PubMed PMID: 18711432.

45. Kopecky J, Kuthejlova M, Pechova J. Salivary gland extract from Ixodes ricinus ticks inhibits production of interferon-gamma by the upregulation of interleukin-10. Parasite Immunol. 1999;21(7):351–6. PubMed PMID: 10417669.

46. Horka H, Staudt V, Klein M, Taube C, Reuter S, Dehzad N, et al. The tick salivary protein sialostatin L inhibits the Th9-derived production of the asthma-promoting cytokine IL-9 and is effective in the prevention of experimental asthma. J Immunol. 2012;188(6):2669–76. doi: 10.4049/jimmunol.1100529. PubMed PMID: 22327077; PubMed Central PMCID: PMCPMC3523721.

47. Preston SG, Majtan J, Kouremenou C, Rysnik O, Burger LF, Cabezas Cruz A, et al. Novel immunomodulators from hard ticks selectively reprogramme human dendritic cell responses. PLoS Pathog. 2013;9(6):e1003450. doi: 10.1371/journal.ppat.1003450. PubMed PMID: 23825947; PubMed Central PMCID: PMCPMC3695081.

48. Salat J, Paesen GC, Rezacova P, Kotsyfakis M, Kovarova Z, Sanda M, et al. Crystal structure and functional characterization of an immunomodulatory salivary cystatin from the soft tick Ornithodoros moubata. Biochem J. 2010;429(1):103–12. doi: 10.1042/BJ20100280. PubMed PMID: 20545626; PubMed Central PMCID: PMCPMC3523712.

49. Coeshott C, Ohnemus C, Pilyavskaya A, Ross S, Wieczorek M, Kroona H, et al. Converting enzyme-independent release of tumor necrosis factor alpha and IL-1beta from a stimulated human monocytic cell line in the presence of activated neutrophils or purified proteinase 3. Proc Natl Acad Sci U S A. 1999;96(11):6261–6. PubMed PMID: 10339575; PubMed Central PMCID: PMCPMC26869.

50. Kwiatkowski K, Mika J. The importance of chemokines in neuropathic pain development and opioid analgesic potency. Pharmacol Rep. 2018;70(4):821–30. doi: 10.1016/j.pharep.2018.01.006. PubMed PMID: 30122168.

51. Turner MD, Nedjai B, Hurst T, Pennington DJ. Cytokines and chemokines: At the crossroads of cell signalling and inflammatory disease. Biochim Biophys Acta. 2014;1843(11):2563–82. doi: 10.1016/j.bbamcr.2014.05.014. PubMed PMID: 24892271.

52. Suffee N, Hlawaty H, Meddahi-Pelle A, Maillard L, Louedec L, Haddad O, et al. RANTES/CCL5-induced pro-angiogenic effects depend on CCR1, CCR5 and glycosaminoglycans. Angiogenesis. 2012;15(4):727–44. doi: 10.1007/s10456-012-9285-x. PubMed PMID: 22752444.

53. Adachi Y, Aoki C, Yoshio-Hoshino N, Takayama K, Curiel DT, Nishimoto N. Interleukin-6 induces both cell growth and VEGF production in malignant mesotheliomas. Int J Cancer. 2006;119(6):1303–11. doi: 10.1002/ijc.22006. PubMed PMID: 16642474.

54. Mark KS, Trickler WJ, Miller DW. Tumor necrosis factor-alpha induces cyclooxygenase-2 expression and prostaglandin release in brain microvessel endothelial cells. J Pharmacol Exp Ther. 2001;297(3):1051–8. PubMed PMID: 11356928.

55. Salcedo R, Young HA, Ponce ML, Ward JM, Kleinman HK, Murphy WJ, et al. Eotaxin (CCL11) induces in vivo angiogenic responses by human CCR3+ endothelial cells. J Immunol. 2001;166(12):7571–8. PubMed PMID: 11390513.

56. Arendse B, Van Snick J, Brombacher F. IL-9 is a susceptibility factor in Leishmania major infection by promoting detrimental Th2/type 2 responses. J Immunol. 2005;174(4):2205–11. PubMed PMID: 15699153.

57. Barron L, Dooms H, Hoyer KK, Kuswanto W, Hofmann J, O’Gorman WE, et al. Cutting edge: mechanisms of IL-2-dependent maintenance of functional regulatory T cells. J Immunol. 2010;185(11):6426–30. doi: 10.4049/jimmunol.0903940. PubMed PMID: 21037099; PubMed Central PMCID: PMCPMC3059533.

58. Chinen T, Kannan AK, Levine AG, Fan X, Klein U, Zheng Y, et al. An essential role for the IL-2 receptor in Treg cell function. Nat Immunol. 2016;17(11):1322–33. doi: 10.1038/ni.3540. PubMed PMID: 27595233; PubMed Central PMCID: PMCPMC5071159.

59. Brake DK, Perez de Leon AA. Immunoregulation of bovine macrophages by factors in the salivary glands of Rhipicephalus microplus. Parasit Vectors. 2012;5:38. doi: 10.1186/1756-3305-5-38. PubMed PMID: 22333193; PubMed Central PMCID: PMCPMC3320552.

60. Takatsu K. Interleukin-5 and IL-5 receptor in health and diseases. Proc Jpn Acad Ser B Phys Biol Sci. 2011;87(8):463–85. PubMed PMID: 21986312; PubMed Central PMCID: PMCPMC3313690.

61. Nuttall PA, Trimnell AR, Kazimirova M, Labuda M. Exposed and concealed antigens as vaccine targets for controlling ticks and tick-borne diseases. Parasite Immunol. 2006;28(4):155–63. doi: 10.1111/j.1365-3024.2006.00806.x. PubMed PMID: 16542317.

62. Kim TK, Tirloni L, Pinto AF, Moresco J, Yates JR, 3rd, da Silva Vaz I, Jr., et al. Ixodes scapularis Tick Saliva Proteins Sequentially Secreted Every 24 h during Blood Feeding. PLoS Negl Trop Dis. 2016;10(1):e0004323. doi: 10.1371/journal.pntd.0004323. PubMed PMID: 26751078; PubMed Central PMCID: PMCPMC4709002.

63. Bonnet S, Jouglin M, Malandrin L, Becker C, Agoulon A, L’Hostis M, et al. Transstadial and transovarial persistence of Babesia divergens DNA in Ixodes ricinus ticks fed on infected blood in a new skin-feeding technique. Parasitology. 2007;134(Pt 2):197–207. doi: 10.1017/S0031182006001545. PubMed PMID: 17076925.

64. Micsonai A, Wien F, Kernya L, Lee YH, Goto Y, Refregiers M, et al. Accurate secondary structure prediction and fold recognition for circular dichroism spectroscopy. Proc Natl Acad Sci U S A. 2015;112(24):E3095–103. doi: 10.1073/pnas.1500851112. PubMed PMID: 26038575; PubMed Central PMCID: PMCPMC4475991.

65. Schuck P. Size-distribution analysis of macromolecules by sedimentation velocity ultracentrifugation and lamm equation modeling. Biophys J. 2000;78(3):1606–19. doi: 10.1016/S0006-3495(00)76713-0. PubMed PMID: 10692345; PubMed Central PMCID: PMCPMC1300758.

66. Reis C, Cote M, Le Rhun D, Lecuelle B, Levin ML, Vayssier-Taussat M, et al. Vector competence of the tick Ixodes ricinus for transmission of Bartonella birtlesii. PLoS Negl Trop Dis. 2011;5(5):e1186. Epub 2011/06/10. doi: 10.1371/journal.pntd.0001186. PubMed PMID: 21655306; PubMed Central PMCID: PMCPMC3104967.

67. Bugarel M, Beutin L, Scheutz F, Loukiadis E, Fach P. Identification of genetic markers for differentiation of Shiga toxin-producing, enteropathogenic, and avirulent strains of Escherichia coli O26. Appl Environ Microbiol. 2011;77(7):2275–81. doi: 10.1128/AEM.02832-10. PubMed PMID: 21317253; PubMed Central PMCID: PMCPMC3067419.

68. Almazan C, Bonnet S, Cote M, Slovak M, Park Y, Simo L. A Versatile Model of Hard Tick Infestation on Laboratory Rabbits. J Vis Exp. 2018;(140). doi: 10.3791/57994. PubMed PMID: 30346382; PubMed Central PMCID: PMCPMC6235438.

69. Patton TG, Dietrich G, Brandt K, Dolan MC, Piesman J, Gilmore RD, Jr. Saliva, salivary gland, and hemolymph collection from Ixodes scapularis ticks. J Vis Exp. 2012;(60). Epub 2012/03/01. doi: 10.3791/3894. PubMed PMID: 22371172; PubMed Central PMCID: PMCPMC3912584.

70. Schmittgen TD, Livak KJ. Analyzing real-time PCR data by the comparative C(T) method. Nat Protoc. 2008;3(6):1101–8. Epub 2008/06/13. PubMed PMID: 18546601.

71. Koci J, Simo L, Park Y. Validation of internal reference genes for real-time quantitative polymerase chain reaction studies in the tick, Ixodes scapularis (Acari: Ixodidae). J Med Entomol. 2013;50(1):79–84. Epub 2013/02/23. PubMed PMID: 23427655; PubMed Central PMCID: PMCPMC3703510.

72. Simo L, Slovak M, Park Y, Zitnan D. Identification of a complex peptidergic neuroendocrine network in the hard tick, Rhipicephalus appendiculatus. Cell Tissue Res. 2009;335(3):639–55. Epub 2008/12/17. doi: 10.1007/s00441-008-0731-4. PubMed PMID: 19082627; PubMed Central PMCID: PMCPMC3573535.

73. Jourdi G, Siguret V, Martin AC, Golmard JL, Godier A, Samama CM, et al. Association rate constants rationalise the pharmacodynamics of apixaban and rivaroxaban. Thromb Haemost. 2015;114(1):78–86. Epub 2015/03/13. doi: 10.1160/TH14-10-0877. PubMed PMID: 25761505.

74. Jourdi G, Gouin-Thibault I, Siguret V, Gandrille S, Gaussem P, Le Bonniec B. FXa-alpha2-Macroglobulin Complex Neutralizes Direct Oral Anticoagulants Targeting FXa In Vitro and In Vivo. Thromb Haemost. 2018;118(9):1535–44. Epub 2018/08/03. doi: 10.1055/s-0038-1667014. PubMed PMID: 30071567.

75. Carreau A, Kieda C, Grillon C. Nitric oxide modulates the expression of endothelial cell adhesion molecules involved in angiogenesis and leukocyte recruitment. Exp Cell Res. 2011;317(1):29–41. doi: 10.1016/j.yexcr.2010.08.011. PubMed PMID: 20813110.

76. Paprocka M, Krawczenko A, Dus D, Kantor A, Carreau A, Grillon C, et al. CD133 positive progenitor endothelial cell lines from human cord blood. Cytometry A. 2011;79(8):594–602. doi: 10.1002/cyto.a.21092. PubMed PMID: 21710642.

77. Carpentier G. Angiogenesis analyser tool for Image J. 2012.

78. Kelley LA, Mezulis S, Yates CM, Wass MN, Sternberg MJ. The Phyre2 web portal for protein modeling, prediction and analysis. Nat Protoc. 2015;10(6):845–58. Epub 2015/05/08. doi: 10.1038/nprot.2015.053. PubMed PMID: 25950237; PubMed Central PMCID: PMCPMC5298202.

